# CHAIR an antigen-resolved antibody-reactome resource for healthy adult immune aging

**DOI:** 10.64898/2026.09.29.755197

**Authors:** Gustav N. Sundell, Minghui Chen, Zigan Sha, Xiangyang Sun, Jiameng Yao, Junjie Liu, Chuansheng Hu, Haoyun Xu, Junbiao Dai, Huan Qi, Mingliang Ma, Jinhua Wu, Yang Li, Shujuan Guo, Jinpiao Lin, Huiming Sheng, Sheng-Ce Tao

## Abstract

Antigen-recognition histories accumulate across adult life, but existing human aging resources largely emphasize host-state measurements rather than antigen-resolved humoral memory. We present CHAIR, the Chinese Healthy Aging Immune Resource, which integrates self-antigen, human virome and toxin/virulence-factor PhIP-seq with routine clinical measurements and T/B/NK phenotyping in clinically screened adults aged 20-94 years. The resource framework includes 1,008 unique PhIP-seq samples, 995 clinical records, 977 T/B/NK records and 853 participants with all three layers. Across modeled spaces comprising 23,584 self-antigen targets, 210 virome targets and 1,258 virulence-factor source targets, most measured antibody-reactivity architecture was consistent with broad preservation within domains, whereas age- and sex-associated signals were concentrated in focal targets and modules. Together, CHAIR provides a traceable, antigen-resolved reference framework based on clinically screened adults for investigating focal humoral remodeling across healthy adult aging.

## Introduction

Human aging resources increasingly move beyond single biomarkers toward cohort-scale views of molecular, cellular and clinical change. Multi-omics and plasma-proteome atlases have shown that aging follows nonlinear trajectories across adult life^1,2^. Chinese and multimodal clock efforts have linked aging phenotypes across population, molecular and clinical layers^3,4^, and proteomic clocks further connect molecular age estimates with mortality and age-related disease risk^5^. In parallel, immune-cell studies have mapped single-cell aging states and sex-dimorphic immune trajectories^6,7^, recent reviews synthesize age-associated immune remodeling^8^, and T, B and NK cell studies define compartment-level changes over adult life^9–12^. Systems-immunology frameworks further emphasize the value of population variation for interpreting human immune phenotypes^13^. These resources have clarified host-state remodeling, but they provide less direct information about antigen-recognition histories that accumulate through infection, vaccination, tissue surveillance and environmental exposure.

Circulating antibodies offer a complementary view because they preserve antigen-specific memory, but conventional serology usually measures a narrow set of pathogens or autoantigens. Phage immunoprecipitation and sequencing (PhIP-seq) enables peptide-resolution profiling across large antigen libraries^14–16^. Population-scale antibody-repertoire studies have shown that epitope repertoires are individualized and comparatively stable in adults^17,18^, shaped by genetic, environmental and intrinsic factors^19^, and associated with viral and microbiome-linked exposure histories that can vary with age and inflammation^20,21^. Self-antigen PhIP-seq and autoantibodyome studies have also shown that measurable self-reactivity can be present in healthy individuals^15,22^, while disease-focused autoreactivity studies underscore the need for careful interpretation before assigning clinical meaning to self-antigen signals^23,24^. Together, these findings support antibody profiling as a useful resource layer for aging research, while setting boundaries: antigen-level reactivity is not by itself evidence of active infection, causal exposure, clinical autoimmunity or disease mechanism.

Here we present the Chinese Healthy Aging Immune Resource (CHAIR), an antigen-resolved antibody-reactome resource for healthy adult immune aging across three complementary antigenic spaces: human self-antigens, the human virome and virulence factors. Together these libraries capture endogenous autoreactivity, common viral immune memory and environmentally acquired microbial/toxin-associated antigen exposure. CHAIR integrates these three antigenic PhIP-seq spaces with routine clinical measurements and T/B/NK immune-cell phenotyping in clinically screened Chinese adults aged 20-94 years. The framework includes 1,008 individual PhIP-seq samples, 995 clinical records, 977 T/B/NK records and 853 participants with all three layers. We use CHAIR to ask whether healthy adult aging is accompanied by broad humoral erosion or expansion, or instead by preserved antibody-reactome architecture with focal remodeling of selected self, viral and virulence-factor targets. The resource value of CHAIR lies in the linked antigen-space matrices, participant-modality bridge, data dictionaries, source traces and access tiers, rather than in any single marker or prediction model. We organize the resulting matrices, model outputs, source traces and access tiers as a traceable Resource for bounded studies of antigen-specific immune aging.

## Results

### CHAIR links antigen-resolved humoral profiles with clinical and immune-cell phenotyping

We developed CHAIR as a clinically screened healthy-adult Resource linking antigen-resolved antibody-reactome profiling with participant-level clinical and cellular immune phenotyping. We first defined the CHAIR data model by connecting the healthy-aging cohort, three PhIP-seq antigen spaces, clinical screening, T/B/NK phenotyping, processed matrices, data dictionaries, source-data products and reproducibility maps through a participant and sample bridge (Fig. 1a).

**Figure 1.**
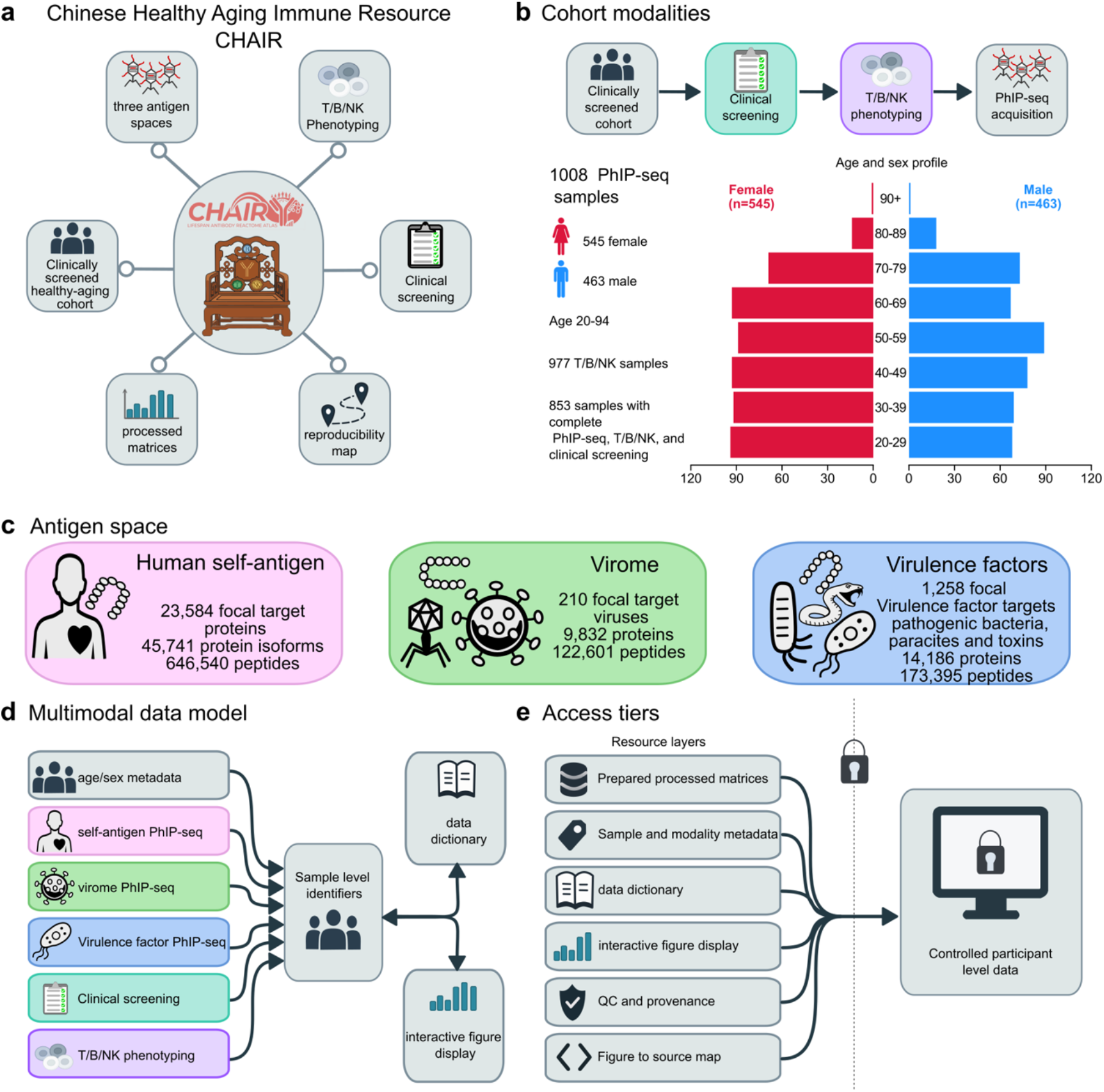
CHAIR resource architecture and healthy-aging cohort design. a) CHAIR is organized as a clinically screened, antigen-resolved immune-aging resource linking baseline PhIP-seq samples, three antigen-space libraries, clinical variables, T/B/NK phenotyping, processed matrices, source-data tables, data dictionaries, and reproducibility maps. The base PhIP-seq resource contains 1,008 deduplicated samples. b) Cohort and modality availability. The base PhIP-seq cohort includes 1,008 samples from participants aged 20-94 years, including 545 female and 463 male participants. Matched resource layers include 995 samples with extensive clinical laboratory screening results, 977 samples with T/B/NK phenotyping data, and 853 records overlapping across the T/B/NK, clinical screening record, and deduplicated PhIP-seq base datasets. Global cohort details are provided in Figure S1 and Table S1. c) Three color-coded PhIP-seq antigen spaces define the antibody-reactome component of CHAIR. The human self-antigen library contains 646,540 peptides mapping to 23,584 focal target proteins. The virome library contains 122,601 peptides grouped into 210 focal target viruses. The virulence-factor library contains 173,395 peptide rows summarized into 1,258 source targets. d) Multimodal data model connecting PhIP-seq layers, clinical measurements, T/B/NK phenotyping, age/sex metadata, data dictionary fields, and figure source-data products through sample-level and modality-level identifiers. e) Resource access and privacy-aware data layers. The current package indexes prepared processed matrices, sample and modality metadata, data dictionaries, interactive figure displays, QC/provenance records, figure-to-source maps, and a CHAIR interface layer undergoing final access and privacy review.

The participant bridge spans 1,205 unique individuals across available study modalities (Supplementary Fig. 1; Supplementary Table 1). The core PhIP-seq cohort included 1,008 adults aged 20–94 years (545 female and 463 male; Fig. 1b), of whom 995 had clinical screening data and 977 had T/B/NK immune-cell phenotyping; all three data layers were available for 853 participants (Supplementary Table 1).

Technical replicates were used during quality control to validate data integration before generating the final participant resource. Subsequent analyses were performed on the validated unique participant cohort.

CHAIR profiles three complementary antigen spaces (Fig. 1c and Supplementary Fig. 2). The human self-antigen library contains 646,540 peptides summarized for target-level modeling across 23,584 self-antigen targets. The human virome library contains 122,601 peptides modeled across 210 virome targets or species groups. The virulence-factor library contains 173,395 peptides modeled across 1,258 source targets. Raw annotation counts, peptide-to-target mappings and modeled denominators are tracked separately because the three libraries differ in design and aggregation. Clinical and T/B/NK variables are documented with completeness, units and access tiers, and the resource manifest links access logic to the analysis layers (Fig. 1d,e; Supplementary Table 2).

This organization allowed each analysis to use an explicit and traceable denominator in the modeled targets. Global antibody-reactome summaries use the 1,008-sample PhIP-seq base, integrated analyses use modality-specific overlaps, and public summary outputs exclude direct participant-level fields unless a deidentified or controlled-access release tier has been approved. Thus, the first layer of CHAIR is a denominator and provenance framework rather than a single discovery plot. Together, we define the denominator, library and access framework used by subsequent analyses.

### Healthy adult aging shows broadly preserved antibody-reactome architecture with sparse focal associations

We next asked whether healthy adult aging was associated with broad remodeling of antibody-reactome architecture or with focal departures from a largely preserved background. Across the 1,008-sample PhIP-seq resource, age-group summaries showed broadly similar global reactivity patterns within each antigen space (Fig. 2a; Supplementary Fig. 3). Because self-antigen panels use sample-level breadth or score summaries whereas virome and virulence-factor panels use target-level positive fractions or score summaries, these views are interpreted within domain rather than as a shared breadth metric across libraries.

**Figure 2.**
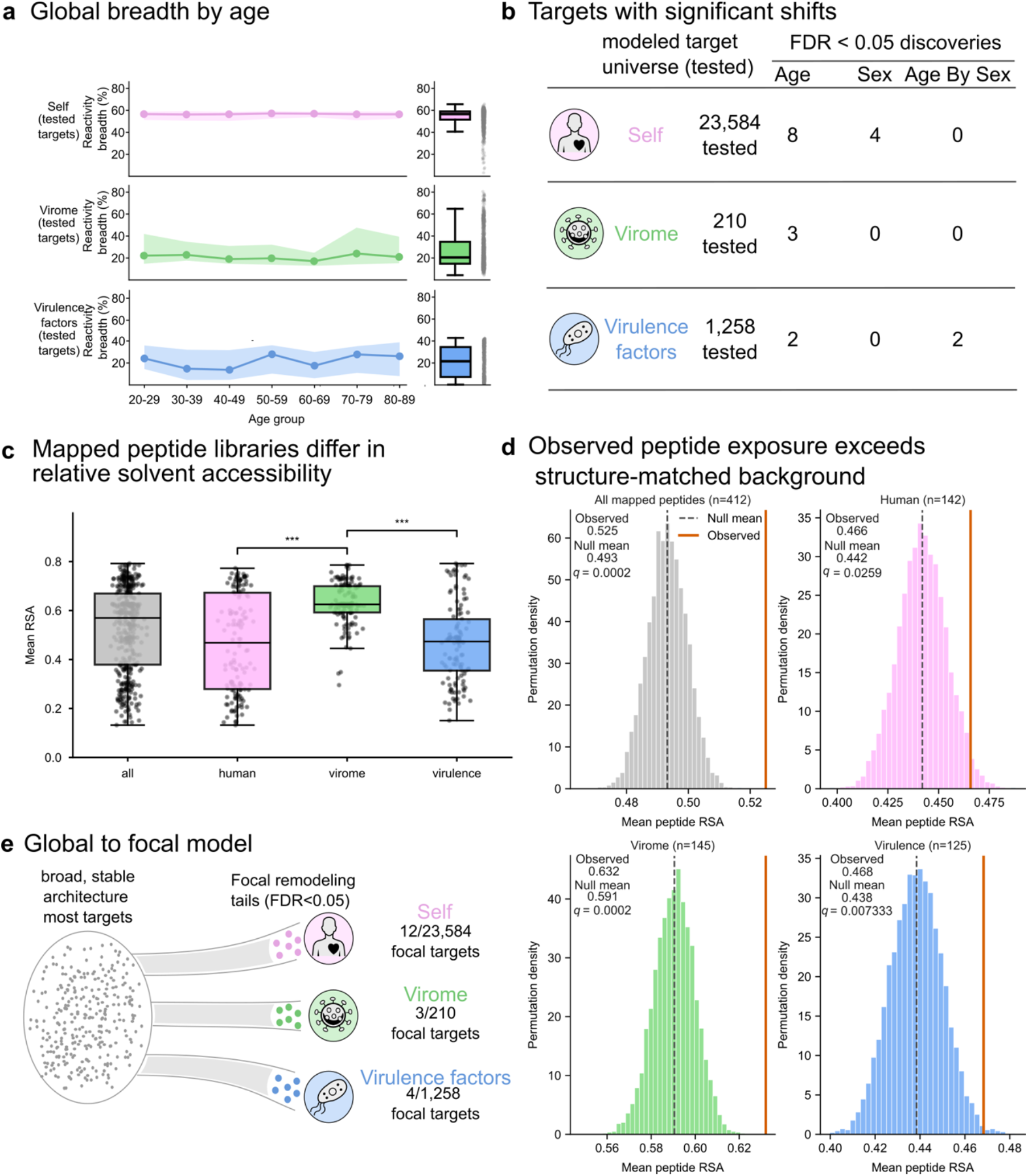
Global stability and focal remodeling across the antibody reactome. a) Metric-specific antibody-reactome summaries across age groups in the 1,008-sample PhIP-seq cohort. Self-antigen panels use sample-level breadth or score summaries, whereas virome and VF panels use target-level positive fraction or score summaries; these metrics are domain-specific architecture summaries rather than measurements based on a shared breadth denominator. b) Model-tested feature spaces included 23,584 self-antigen features, 210 virome targets, and 1,258 VF targets. At q < 0.05, self-antigen results included 8 age-associated and 4 sex-associated targets, with no age-by-sex interactions; virome results included 3 age-associated targets, with no sex or interaction targets; VF results included 2 age-associated and 2 age-by-sex interaction targets, with no sex-associated targets. c) Relative solvent accessibility (RSA) of mapped peptides across human self-antigen, virome, and virulence-factor libraries. Peptides were selected from the top sample-positive peptides and mapped to AlphaFold-derived structures. Strict mapping required ≥85% sequence identity and ≥95% coverage; the relaxed virome approach used homologous structures meeting ≥80% identity and ≥80% coverage. The full mapped dataset comprised 412 peptides (human n = 142, virome n = 145, virulence-factor n = 125). Boxplots show peptide-level mean RSA; points represent individual peptides. Pairwise comparisons used two-sided Mann–Whitney U tests with Holm correction (*** Holm-adjusted *p* < 0.001). d) Structure-matched permutation analysis tested whether mapped peptides occupy more solvent-accessible regions than expected by chance within source proteins. For each peptide, 10,000 random contiguous length-matched windows were sampled from the same structure. Orange lines indicate observed means; dashed lines indicate null means. Empirical P values were corrected using Benjamini–Hochberg FDR across primary tests. e) Selected top focal targets ranked by FDR-supported signal class.

We modeled associations between age, sex and age-by-sex interaction across the self-antigen, virome and virulence-factor antigen spaces, applying false discovery rate (FDR) correction within each target set. FDR-supported target-level signals were sparse relative to the modeled antigen spaces (Fig. 2b; Supplementary Fig. 4; Supplementary Tables 5 and 6). Among 23,584 modeled self-antigen targets, 8 were age-associated and 4 were sex-associated at FDR < 0.05, with no age-by-sex interaction target passing this threshold. Among 210 modeled virome target/species-group units, 3 were age-associated and none showed sex or age-by-sex interaction signals. Among 1,258 modeled virulence-factor source targets, 2 were age-associated and 2 showed age-by-sex interaction signals, whereas no target showed a main sex effect. Combining age, sex and interaction terms yielded focal signals in 12 of 23,584 self-antigen targets (0.051%), 3 of 210 virome units (1.43%) and 4 of 1,258 virulence-factor targets (0.318%). These results do not support global antibody-reactome collapse or expansion across healthy adulthood; statistically supported remodeling was concentrated in small target tails.

We used structural mapping to provide biophysical context for peptides selected from the highest sample-positivity candidates in each antigen space, not to infer prior antigen encounter or binding mechanism. The full mapped peptide set contained 412 peptides, including 142 human self-antigen peptides, 145 virome peptides and 125 virulence-factor peptides. Mean relative solvent accessibility (RSA) was calculated for each peptide. The mean RSA was 0.525, well above the 0.2 threshold generally considered to be solvent accessible^25^. Peptides from the virome library had a higher solvent accessibility than the peptides from self-antigen and virulence factor libraries (Fig. 2c; Supplementary Table 5). Because some virome peptides lacked direct structure matches, the primary set included a virome-only relaxed homologous mapping layer; a strict-only sensitivity set containing 314 directly matched peptides is reported in Supplementary Table 5.

Structure-matched permutation analysis tested whether mapped peptides occupied more solvent-accessible regions than expected from same-protein, same-length random windows (Fig. 2d; Supplementary Table 5). Across all 412 mapped peptides, observed mean relative solvent accessibility was 0.525 compared with a null mean of 0.493 after 10,000 permutations (FDR-adjusted q = 0.0002), with the same direction within each antigen space. Together, these data support a global-to-focal model in which broad summaries remain largely preserved while focal target tails carry age-, sex-or interaction-associated structure (Fig. 2e).

Surface accessibility provides an orthogonal plausibility check that highly reactive peptides preferentially map to exposed regions likely to be antibody accessible. The structural analysis also served as a guardrail for interpretation. Surface accessibility enrichment makes the selected peptide set more plausible as a displayed-antigen reactivity layer, but the analysis remains peptide- and structure-contextual rather than mechanistic. We therefore used it to support traceable annotation of the mapped high-sample-positivity peptide set, while leaving exposure history, epitope conformation and binding validation to future experimental studies. As a Resource layer, these outputs provide reusable global summaries, FDR-supported target landscapes and structural-context annotations while keeping peptide accessibility distinct from mechanistic proof.

### Focal age-associated targets define antigen-space remodeling

We next localized the age-associated component of this global-to-focal pattern using harmonized target-level models across the three CHAIR antigen spaces (Fig. 3a; Supplementary Table 3). The analysis used the 1,008-sample PhIP-seq base set and evaluated 25,052 modeled targets: 23,584 human self-antigen targets, 210 virome target/species-group units and 1,258 virulence-factor source targets. Model outputs were organized by age, sex and age-by-sex terms.

**Figure 3.**
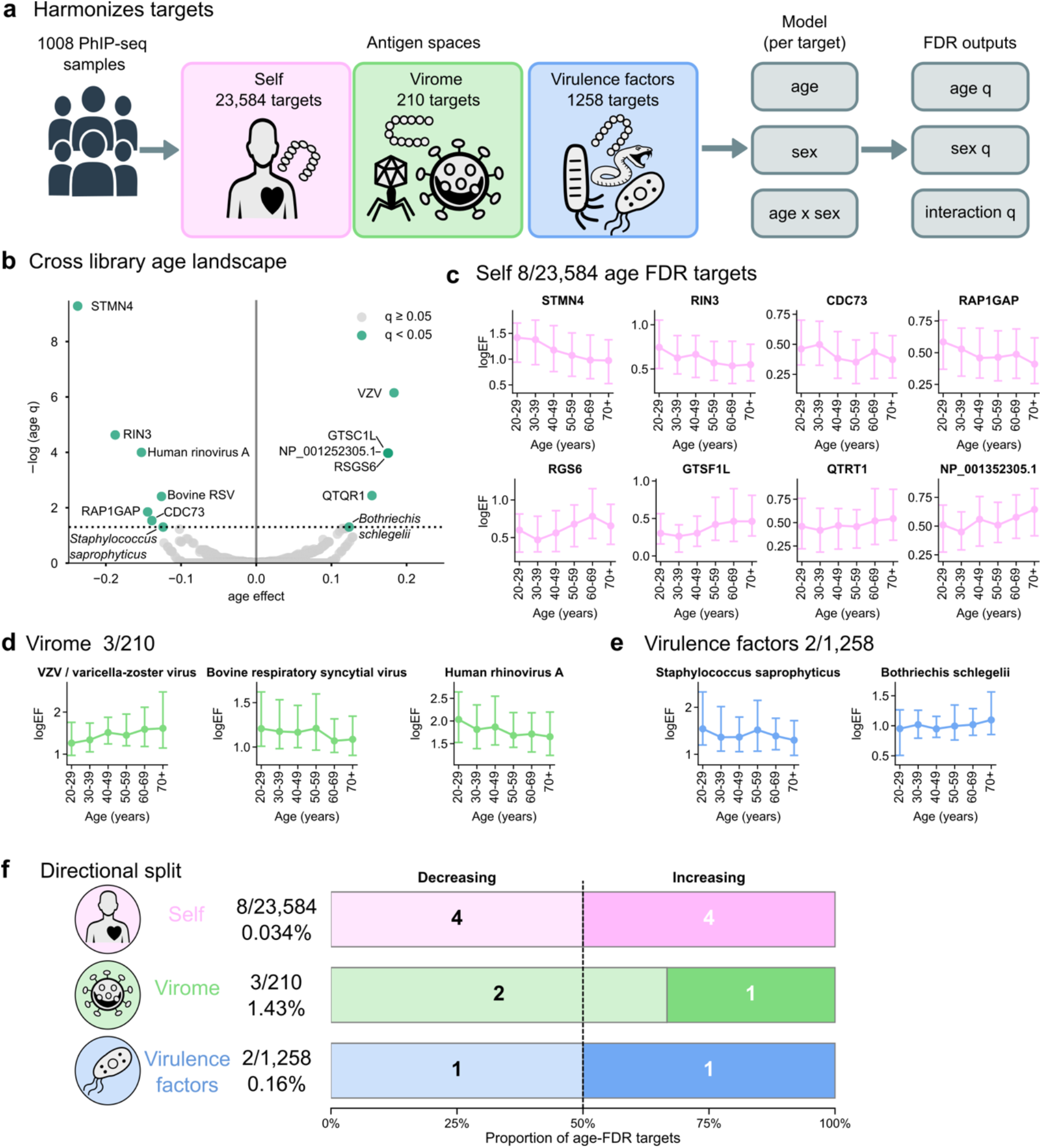
Harmonized target-level age remodeling across self-antigen, virome, and Virulence factor antigen spaces. a) Unified target-level modeling framework applied to the CHAIR PhIP-seq sample set. The analysis uses 1,008 samples and evaluates three modeled target spaces: 23,584 human self-proteome targets, 210 human virome targets, and 1,258 toxin or virulence-factor targets. Model outputs include age, sex, and age-by-sex interaction terms where available, with FDR-adjusted *q* values. Color coding is consistent across figures, with self-antigen targets shown in pink, virome targets in green, and virulence-factor targets in blue. b) Cross-library age-effect landscape across 25,052 modeled antigen targets. Highlighted targets represent age-associated discoveries passing the FDR threshold (q < 0.05). Positive and negative age effects indicate increasing or decreasing target-level antibody-reactivity associations with age, respectively. The plot shows focal remodeling rather than broad domain-wide shifts in a small number of targets. c) Self-antigen age modules. Eight self-antigen targets pass the age-FDR threshold among 23,584 modeled self-antigen targets, with four showing increasing and four showing decreasing age-associated patterns. Self-antigen target displays use fixed Top10 peptide-panel scores. d) Virome age modules. Three virome targets pass the age-FDR threshold among 210 modeled virome targets: varicella-zoster virus (VZV), human rhinovirus A, and bovine respiratory syncytial virus. These seroreactivity patterns are compatible with prior immune recognition and should not be interpreted as evidence of active infection. e) VF age modules. Two virulence-factor targets pass the age-FDR threshold among 1,258 modeled VF targets: Bothriechis schlegelii and Staphylococcus saprophyticus. These represent exploratory antibody-reactivity associations and do not establish validated exposure histories or causal relationships. For panels c–e, points represent age-group summary values, and error bars indicate the interquartile range. f) Age-FDR counts are shown relative to their library-specific tested target spaces: self-antigen 8/23,584, virome 3/210, and VF 2/1,258. Direction splits are self 4 decreasing and 4 increasing, virome 2 decreasing and 1 increasing, and VF 1 increasing and 1 decreasing. Exact effect sizes, q values, peptide-panel definitions, age-bin summaries, and source-table links are provided in the Figure 3 source-data workbook and Tables S4-S7.

Across the combined target landscape, 13 targets passed the age FDR threshold (Fig. 3b). These included 8 of 23,584 self-antigen targets, 3 of 210 virome target/species-group units and 2 of 1,258 virulence-factor source targets. The self-antigen layer contained a balanced directional split, with four age-increasing and four age-decreasing targets (Fig. 3c,f). STMN4 showed the largest absolute self-antigen age coefficient and decreased with age (age effect = −0.238, q = 5.21 × 10⁻¹⁰), followed by RIN3 (age effect = −0.188, q = 2.35 × 10⁻⁵). Age-increasing self-antigen targets included GTSF1L, RGS6 and QTRT1. These patterns are best interpreted as focal target-level shifts in antibody-reactivity magnitude or immunodominance, not as evidence for clinical autoimmunity or broad self-reactivity expansion.

The virome target-level model identified three age-associated target/species-group units (Fig. 3d; Supplementary Table 3). Varicella-zoster virus (VZV) showed an age-increasing association (age effect = 0.183, q = 7.09 × 10⁻⁷), whereas human rhinovirus A and a bovine respiratory syncytial virus cross-reactive/low-prior virome unit showed age-decreasing target-level summaries (age effects = −0.153, q = 1.00 × 10⁻⁴, and −0.126, q = 0.00391, respectively). The virulence-factor layer contained two age-FDR targets among 1,258 modeled source targets (Fig. 3e; Supplementary Fig. 5; Supplementary Table 3): *Bothriechis schlegelii* increased with age (age effect = 0.123, q = 0.0498), whereas *Staphylococcus saprophyticus* decreased (age effect = −0.124, q = 0.0498). These viral and virulence-factor entries are seroreactivity associations, not evidence of active infection, viral load, recent exposure, toxin exposure or causal viral involvement in aging. The target-level and peptide-level source traces support moving from sparse global counts to specific targets, effect directions, peptide-panel definitions and positivity summaries (Supplementary Tables 3 and 4).

### CHAIR identifies sex-aware herpesvirus and DAZ reactivity modules

We next asked whether the focal remodeling identified by target-level models contained sex-aware structure across the CHAIR antigen spaces (Fig. 4; *q* values are provided for these focal targets in Supplementary Table 6). In the harmonized target scan, sex-associated signals were concentrated in the self-antigen layer rather than distributed broadly across all libraries. Four of 23,584 modeled self-antigen targets passed the sex FDR threshold: DAZ2, DAZ3 and DAZ4 were higher in males, whereas ZNF606 was higher in females (Fig. 4a). No virome target/species-group unit and no virulence-factor source target passed the main sex-effect FDR threshold. The virulence-factor layer instead contributed two age-by-sex interaction targets, *Synechococcus lacustris* and *Tropidechis carinatus*, each with interaction q = 0.031 (Fig. 4b). These interaction entries identify age-stratified seroreactivity patterns that differ by sex, not sex-specific toxin, pathogen or environmental exposure histories.

**Figure 4.**
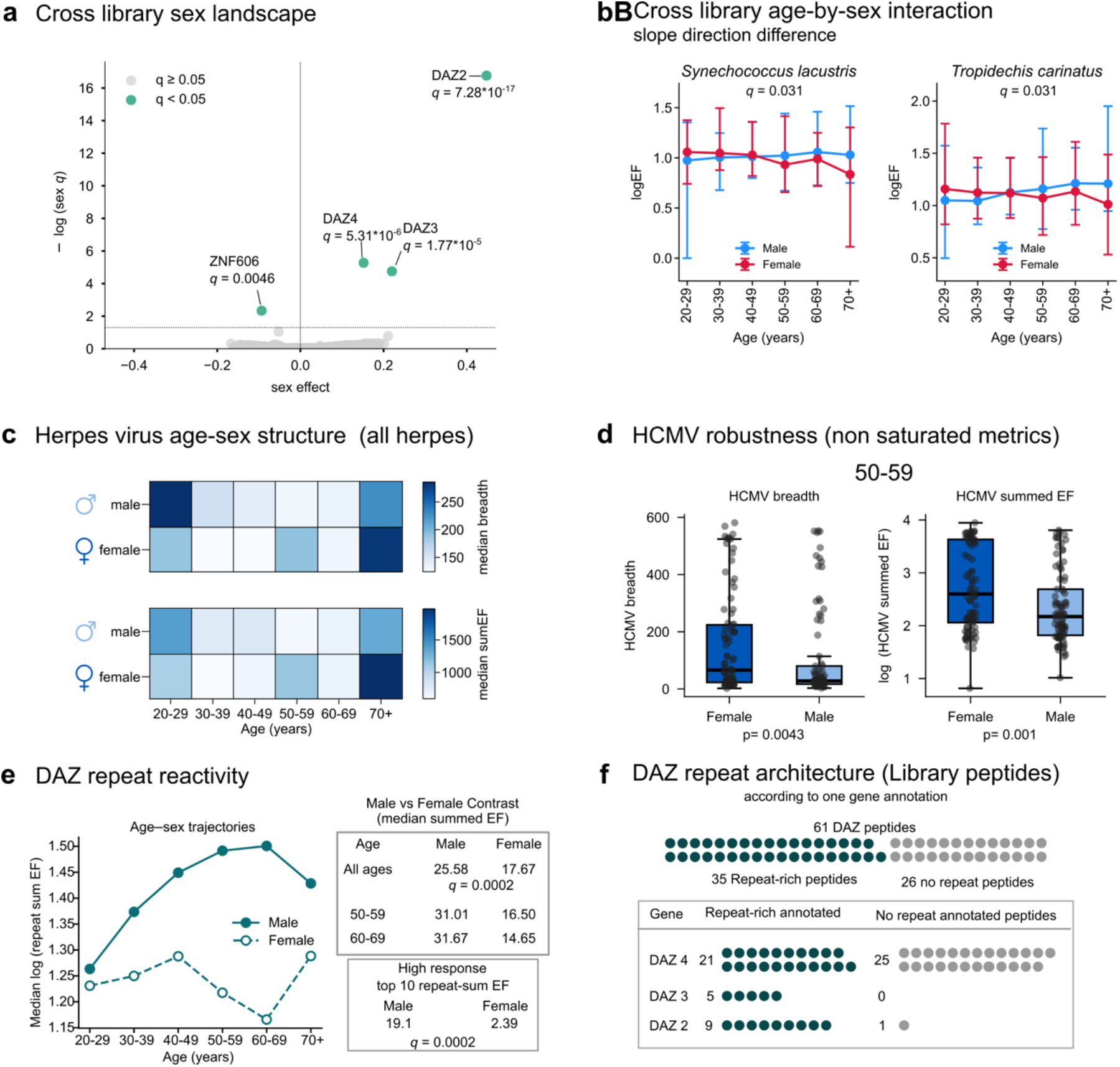
Focal sex-associated DAZ and herpesvirus antibody-reactivity modules. a) Four human self-antigen targets showed sex-associated antibody-reactivity patterns. Three male-enriched targets mapped to DAZ2, DAZ3, and DAZ4, whereas one female-enriched target mapped to ZNF606. b) Two VF targets show age-by-sex interactions in which the association changes with age in opposite directions between females and males. c) Herpesvirus age-sex structure summarized using non-saturated seroreactivity metrics. The heat maps use ALL_HUMAN_HERPES breadth and maximum EF across the 1,008 samples, with stratum-specific sample counts provided in the source data. d) HCMV/CMV robustness analysis. In the female-versus-male HCMV contrast for ages 50-59, median breadth values were 67.00 versus 28.00, and median summed EF values were 418.06 versus 148.70. Corresponding permutation *P* values were 0.004 and 0.001, respectively. CMV strict positivity was saturated in this contrast and should be interpreted only as context. e) DAZ repeat-reactivity magnitude across sex-stratified age groups. Across all samples, median DAZ repeat-sum EF values were 25.58 in males versus 17.67 in females, with a permutation-adjusted q value of 0.0002. Midlife contrasts showed repeat-sum EF medians of 31.01 vs 16.50 in the 50-59 stratum and 31.67 vs 14.65 in the 60-69 stratum. f) DAZ repeat architecture. The repeat metadata table contains 61 DAZ peptides annotated across three of the four DAZ proteins, including 35 repeat-rich peptides and 26 non-repeat-rich peptides. Repeat-rich peptide counts were 21 for DAZ4, 5 for DAZ3, and 9 for DAZ2. As seen in Figure S7, the DAZ proteins are highly homologous, which means that peptides annotated to one DAZ homologue often map to additional DAZ homologues because of sequence similarity within this protein family.

Because binary herpesvirus positivity was nearly saturated in this cohort, we summarized herpesvirus structure using non-saturated breadth and summed-EF metrics rather than positive/negative calls (Fig. 4c,d; Supplementary Fig. 6; Supplementary Table 6). Across annotated human herpesvirus peptides, age-by-sex strata showed visible structure in median breadth and summed EF. A focused HCMV robustness analysis highlighted the 50-59-year stratum, where female and male sample counts were balanced (n = 89 each). In this stratum, females had higher median HCMV breadth than males (67.00 versus 28.00 positive peptides; permutation P = 0.004) and higher median HCMV summed EF (418.06 versus 148.70; permutation P = 0.001). These patterns are seroreactivity breadth and magnitude summaries, not active infection, viral load, recent exposure, individual infection status or a menopause mechanism.

The self-antigen sex signal was dominated by a DAZ repeat-reactivity module (Fig. 4e,f; Supplementary Fig. 7). Across all 1,008 PhIP-seq samples, median DAZ repeat-sum EF was higher in males than females (25.58 versus 17.67; permutation-adjusted q = 0.0002). The same direction was observed in the 50-59-year stratum (31.01 versus 16.50) and the 60-69-year stratum (31.67 versus 14.65). High-response enrichment showed a similar pattern: 19.01% of males versus 2.39% of females fell into the top 10% of DAZ repeat-sum EF values. Repeat metadata mapped 61 DAZ peptides across DAZ2, DAZ3 and DAZ4, including 35 repeat-rich and 26 non-repeat-rich peptides. Because DAZ paralogues are highly homologous, the module is interpreted as repeat-epitope antibody reactivity in a subset of men, not as a male-exclusive marker, infertility biomarker, causal reproductive autoimmunity result or blood-testis-barrier mechanism.

### Multidomain antigen profiling enables mimotope bridge prioritization

CHAIR profiles self, viral and toxin/virulence-factor antigen spaces in the same PhIP-seq sample set, enabling cross-domain analyses that are not available from a single antigen library alone. We used this structure to build a homology-aware bridge workflow for nominating self-foreign peptide pairs with short sequence similarity and aggregate co-reactivity support (Fig. 5a; Supplementary Figs. 8 and 9; Supplementary Table 7). The workflow began from candidate atlases filtered by sample positivity: 160,072 human peptides were retained from the self-antigen atlas, and 20,295 viral or toxin/virulence-factor peptides were retained from the foreign-antigen atlases. We then searched for shared 7-amino-acid seed motifs, evaluated 24,501 sequence pairs and retained 20,000 ranked bridge candidate pairs for reporting. Candidate scoring combined exact shared span, biochemical similarity span, Pearson correlation of logEF values, positive-overlap enrichment, both-positive sample counts and homology-risk filters. The purpose was candidate prioritization, not validation of molecular mimicry or infection-induced autoimmunity. The resulting table is therefore organized as a reusable evidence ladder whose rows can be filtered by sequence span, co-reactivity support, homology risk and source provenance.

**Figure 5.**
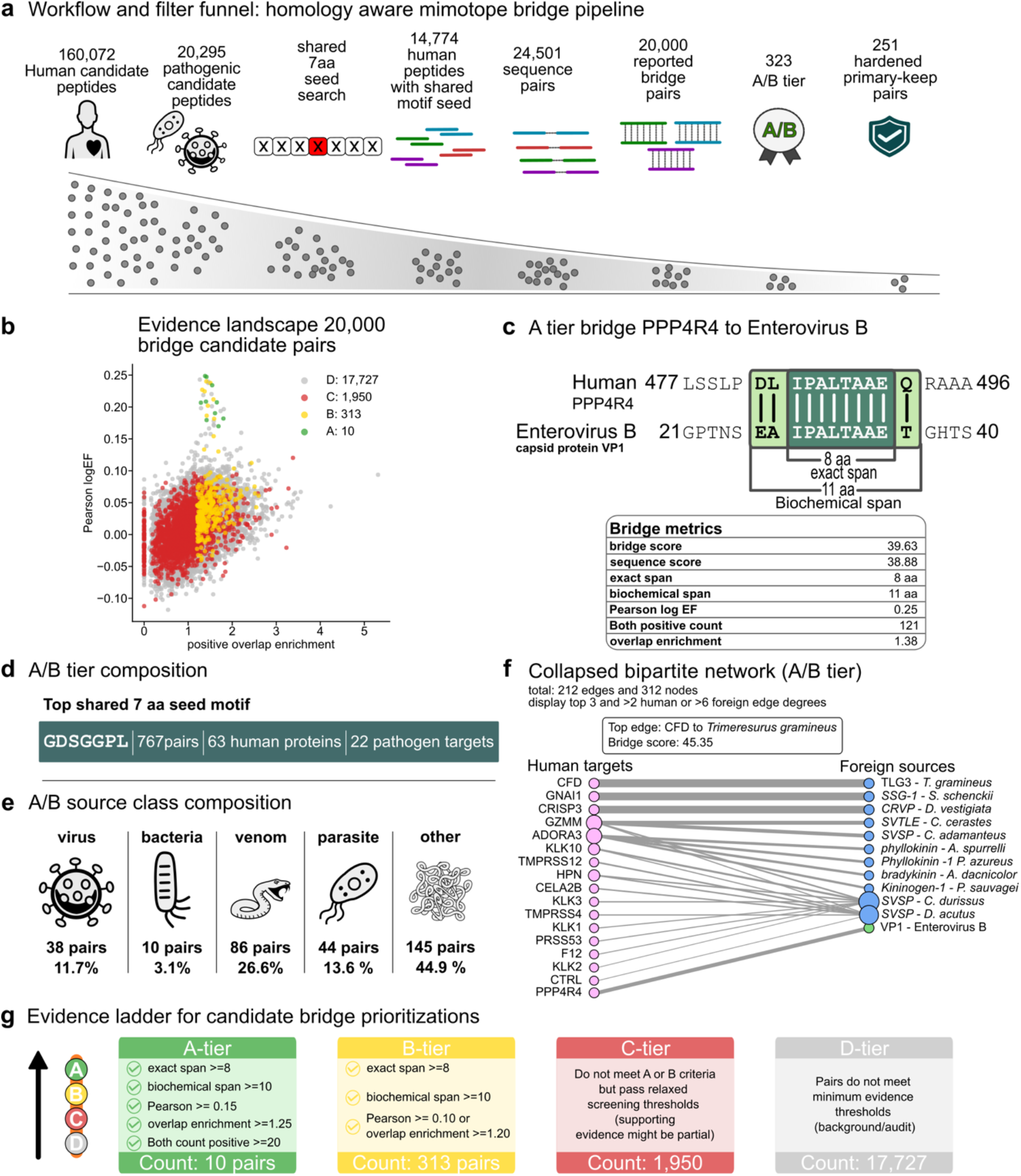
Homology-aware mimotope bridge discovery and prioritization framework. a) Workflow and filter funnel for self-foreign mimotope bridge nomination. The analysis begins with 160,072 human candidate peptides and 20,295 pathogen/toxin candidate peptides, identifies shared 7-aa seed motifs, evaluates 24,501 sequence pairs, and generates 20,000 ranked bridge candidate pairs. A/B-tier candidates included 323 pairs, of which 251 remain in the hardened primary-keep set. b) Evidence landscape across 20,000 ranked bridge candidate pairs. Candidate counts by evidence tier were A = 10, B = 313, C = 1,950, and D = 17,727. A-tier candidates had a median bridge score of 37.48, a median exact span of 8.0 aa, a median biochemical span of 11.0 aa, a median Pearson correlation of logEF values of 0.207, and a median positive-overlap enrichment of 1.48. c) Representative A-tier candidate bridge linking PPP4R4 and Enterovirus B. The top-ranked representative candidate has a bridge score of 39.63, exact span 8, biochemical span 11, Pearson logEF 0.249, both-positive count of 121 and a positive-overlap enrichment of 1.38. d) Shared motif and source-class landscape. The most frequent shared 7-aa seed motif is GDSGGPL, which appears in 767 bridge pairs, 63 human proteins, and 22 pathogen targets. e) Retained A/B-tier candidates span five source classes: virus, bacteria, venom, parasite, and other. f) Collapsed human-target and foreign-source candidate network. The total network contains 212 collapsed edges and 312 nodes. The highest-scoring displayed edge connects CFD with TLG3 protein from *T. gramineus*, with a maximum bridge score of 45.35. g) Evidence ladder for follow-up prioritization. A-tier candidates were defined by the following criteria: exact span ≥8 aa, biochemical span ≥10 aa, Pearson logEF ≥0.15, positive-overlap enrichment ≥1.25, both-positive count ≥20, and absence of low-complexity or long-homology flags. B-tier candidates require an exact span ≥8 aa and a biochemical span ≥10 aa, together with either Pearson logEF ≥0.10 or positive-overlap enrichment ≥1.20.

The bridge landscape was tiered to separate higher-priority hypotheses from sequence-only or audit-background matches (Fig. 5b,g; Supplementary Table 7). Among 20,000 ranked pairs, 10 were assigned to tier A, 313 to tier B, 1,950 to tier C and 17,727 to tier D. A/B-tier evidence comprised 323 pairs, of which 251 remained in the hardened primary-keep set after secondary review and additional homology and repeat-like motif filters. The highest-priority displayed A-tier example connected PPP4R4 with Enterovirus B, sharing the IPALTAAE exact span, with bridge score 39.63, exact span 8, biochemical span 11, Pearson logEF 0.249, both-positive count 121 and positive-overlap enrichment 1.38 (Fig. 5c).

We further summarized the candidate table through motif and network views that retain the evidence ladder for follow-up prioritization (Fig. 5d-f; Supplementary Table 7). The most frequent shared 7-amino-acid seed motif was GDSGGPL, observed in 767 bridge pairs, 63 human proteins and 22 foreign targets. A/B-tier source-class composition spanned viral, bacterial or microbial virulence-factor, venom or toxin, eukaryotic or parasite-like and other toxin/virulence-factor sources. A collapsed bipartite network contained 212 edges and 312 nodes, with the highest-scoring displayed edge connecting CFD to *Trimeresurus gramineus*. We use these views to move from the full bridge table to focused hypotheses while preserving the requirement for orthogonal follow-up.

Because all bridge candidates are linked back to peptide identifiers, source classes, co-reactivity scores and homology-risk annotations, this module is useful as a prioritization index even when individual pairs are not taken forward experimentally. It provides a reproducible way to triage a large self-foreign search space into candidates that can be reviewed, filtered, replicated or tested with independent binding and perturbation assays.

### Multimodal CHAIR features support cohort-specific immune-age modeling

We next used CHAIR to ask whether antigen-resolved antibody reactivity could be integrated with cellular and clinical measurements into a compact immune-aging summary. This analysis was designed as a resource use case rather than as a clinical prediction tool. We therefore constructed a cohort-specific Immune Age Index (IAI) with two complementary layers: an interpretable compact complete-case model and a broader near-full internal validation workflow (Fig. 6a; Supplementary Fig. 10; Supplementary Table 8).

**Figure 6.**
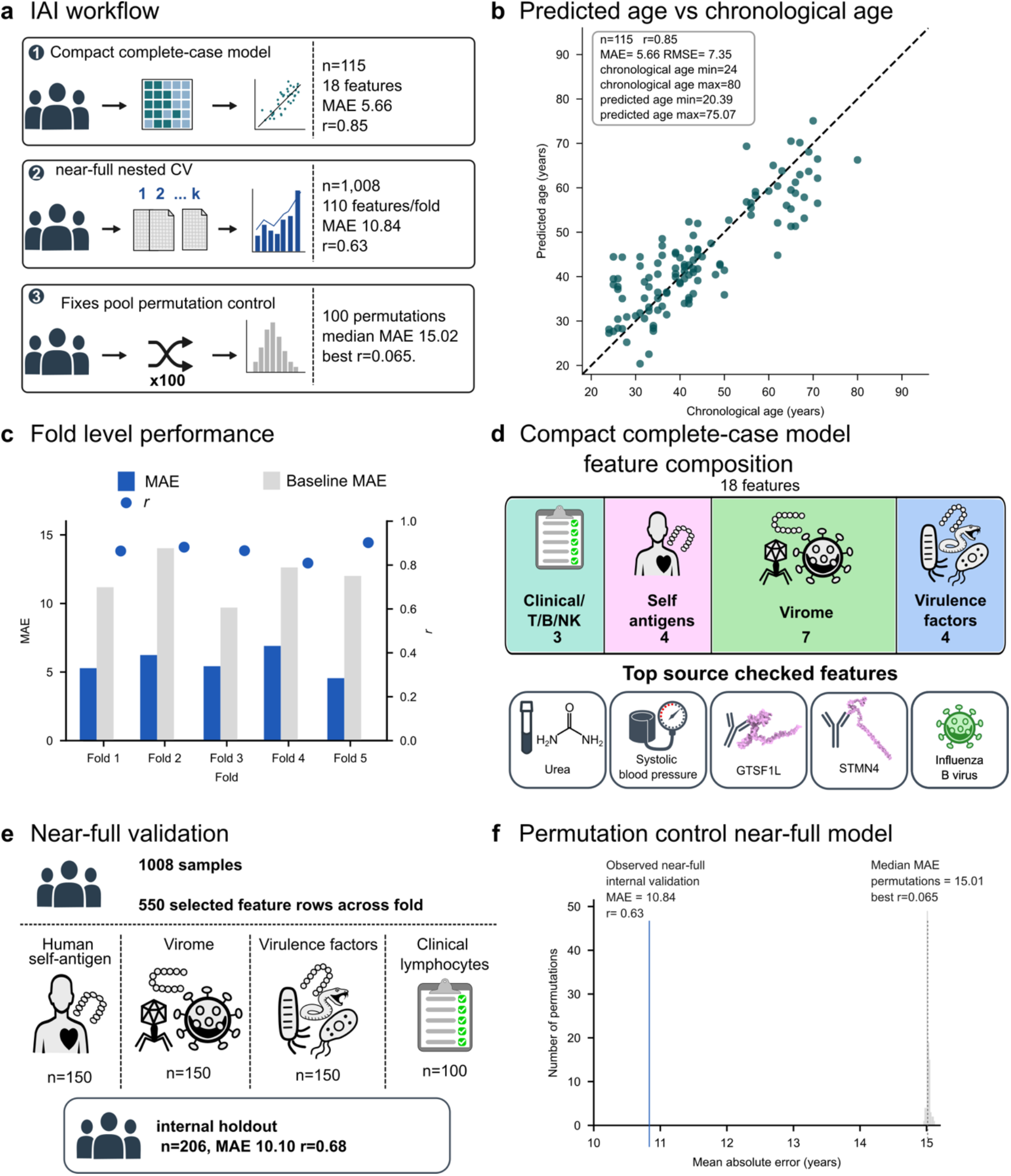
Cohort-specific immune age index derived from multimodal immune features. a) IAI model workflow. IAI represents a model-derived estimate of chronological age from multimodal immune features. The compact complete-case model uses 115 samples and 18 features, with internal cross-validated MAE 5.66, RMSE 7.35, and r 0.85. Near-full nested cross-validation uses 1,008 samples and 110 features per fold, with MAE = 10.84, RMSE 13.49, Pearson correlation between predicted and chronological age (r = 0.63). Fixed-pool permutation controls include 100 permutations. b) Compact-model predicted versus chronological age for 115 complete-case predictions. The scatter plot uses cross-validated predictions from the fixed 18-feature ElasticNet model. c) Compact-model cross-validation performance across five folds. Fold-level MAE ranged from 4.56 to 6.90 years, and Pearson correlation ranged from 0.81 to 0.90. d) The fixed 18-feature panel includes three clinical/lymphocyte features, four human self-antigen PhIP-seq features, seven virome PhIP-seq features, and four VF PhIP-seq features. Feature labels and feature-level correlations are provided in supplementary table S10. e) Near-full internal validation. The near-full nested CV includes 1,008 samples, 545 female and 463 male, and 110 selected features per fold. The nested cross-validation holdout layer used 802 training samples and 206 held-out samples, with holdout MAE 10.10 years Pearson correlation of 0.68. Across folds, selected-feature rows include human_lib 150, VF_lib 150, virus_lib 150, and clinical_tests 100. f) Permutation control near-full model. Across 100 outcome permutations, median MAE was 15.01 years and the maximum observed Pearson correlation was 0.064, compared to the internal validation of the near-full model of MAE 10.84 and Pearson correlation of 0.63. This control evaluates calibration after selected feature-pool construction and is not a full re-screening permutation pipeline.

The compact model used 115 complete-case samples with an 18-feature ElasticNet panel spanning clinical or lymphocyte measurements, self-antigen PhIP-seq, virome PhIP-seq and toxin or virulence-factor features (Fig. 6b-d). In cross-validation, this model achieved mean absolute error (MAE) of 5.66 years, root mean squared error (RMSE) of 7.35 years and predicted-versus-chronological age correlation r = 0.85. Fold-level performance was consistent across the five compact-model folds, with MAE values from 4.56 to 6.90 years and correlations from r = 0.81 to r = 0.90. The fixed 18-feature panel included three clinical or lymphocyte features, four self-antigen features, seven virome features and four toxin or virulence-factor features. These features describe multimodal model composition, not causal antigen exposure or disease mechanisms.

To test whether age-associated structure persisted beyond the complete-case subset, we applied a near-full nested cross-validation workflow across 1,008 participants (Fig. 6e; Supplementary Figs. 10 and 11). This analysis performed fold-local feature screening over human self-antigen, virome, toxin or virulence-factor, clinical and T/B/NK feature spaces, selecting 110 features per fold. The near-full nested cross-validation summary yielded MAE = 10.84 years, RMSE = 13.49 years and r = 0.63, improving over a baseline MAE of 15.00 years by 4.17 years. A separate internal holdout layer used 802 training samples and 206 holdout samples from age-and sex-stratified folds, yielding holdout MAE = 10.10 years, RMSE = 12.88 years and r = 0.68. Across 100 fixed selected-feature-pool age-label permutations, the median permutation MAE was 15.01 years and the best permutation correlation was r = 0.065 (Fig. 6f). The IAI therefore summarizes internally cross-validated, multimodal age-informative structure within CHAIR, but it is not an externally validated biological-age clock, clinical diagnostic, clinical biomarker or generalizable predictor.

### CHAIR enables queryable antigen-resolved exploration of healthy immune aging

Having described CHAIR-derived use cases across antigen-space architecture, target-level age associations, sex-stratified modules, mimotope candidates and immune-age modeling, we organized the dataset as a queryable, traceable Resource package rather than a static set of figures (Fig. 7); Supplementary Fig. 12 provides the corresponding interface architecture and query workflow. The query model defines entry points by sample, modality, peptide, target, age or sex stratum, claim boundary and figure/source-data reproduction. These entry points connect the 1,008-sample PhIP-seq base with self-antigen, virome and toxin or virulence-factor matrices, clinical screening, T/B/NK phenotyping, mimotope bridge tables, data-dictionary fields and figure source-data records. The resource manifest separates public-summary outputs from deidentified-matrix candidates, controlled participant-level data and controlled identifier-linked files so that reuse can be matched to privacy and provenance requirements.

**Figure 7.**
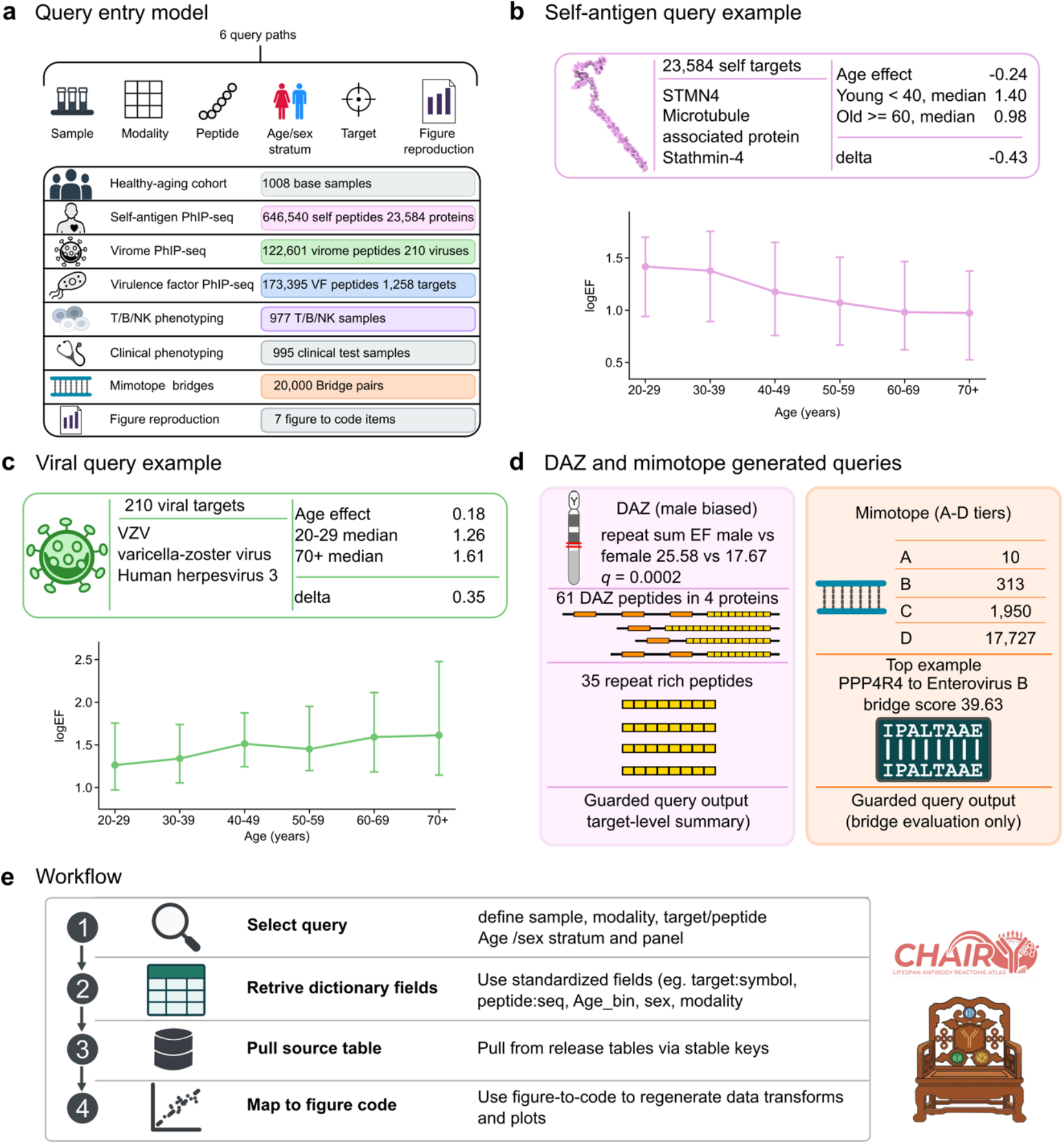
CHAIR as a reusable immune-aging atlas. a) CHAIR query entry framework. The resource manifest defines six query paths and seven modality categories spanning sample metadata, self-antigen PhIP-seq, virome PhIP-seq, toxin/VF PhIP-seq, clinical information, T/B/NK phenotyping, and the CHAIR data dictionary. The base sample count is 1,008. b) Example self-antigen query. The STMN4 target query uses 1,008 samples and the 23,584-target modeled self-antigen feature space. The STMN4 age association coefficient is −0.24; The median score was 1.40 in participants <40 years and 0.98 in participants ≥60 years, and old-minus-young median difference is −0.43. c) Example virome query. The VZV (varicella-zoster virus) query uses the 210-target modeled virome feature space. The VZV age association coefficient is 0.18 with an FDR-adjusted q value of 7.09 × 10⁻⁷. Median score is 1.26 in the 20-29 age bin and 1.61 in the 70+ age group. These represent seroreactivity summaries and should not be interpreted as active infection or viral-load measurements. d) Examples of high-interest queries with interpretation safeguards. The DAZ repeat-sum EF query shows median values of 25.58 in males and 17.67 in females (q = 0.0002); the DAZ architecture table contains 61 peptides, including 35 repeat-rich and 26 non-repeat-rich peptides. The mimotope ladder contains A-tier (10), B-tier (313), C-tier (1,950), and D-tier (17,727) candidate pairs; the top displayed candidate is the PPP4R4–Enterovirus B bridge with a bridge score of 39.63. e) Query workflow illustrating progression from resource selection to modality-specific exploration, target-level interrogation, and figure/source-data retrieval.

We used STMN4 and VZV to demonstrate how a query moves from an entry point to a source-backed target-level result (Fig. 7b,c). In the self-antigen space, STMN4 used the 1,008-sample cohort and 23,584 tested self-targets and showed an age-decreasing association (age effect = −0.238), with median fixed-panel scores lower in participants aged 60 years or older than in those younger than 40 years (0.977 versus 1.404; delta = −0.427). In the virome space, VZV used the 210-target virome model space and showed an age-increasing association (age effect = 0.183; q = 7.09 x 10-7), with displayed median scores increasing from 1.26 in participants aged 20-29 years to 1.61 in participants aged 70 years or older. These outputs are seroreactivity summaries; they do not establish active infection, viral load, exposure timing or a clinical diagnosis.

High-interest guarded queries apply the same structure to DAZ and mimotope examples (Fig. 7d,e). A DAZ repeat-reactivity query links sex-stratified signal, repeat annotation and peptide architecture: the DAZ repeat-sum enrichment score was higher in male than female participants (median 25.58 versus 17.67; q = 0.0002), and the DAZ architecture table indexes 61 DAZ peptides, including 35 repeat-rich and 26 non-repeat-rich peptides. A mimotope query returns bridge candidates rather than validated molecular mimicry events: 20,000 screened bridge pairs were stratified into tiers A-D, including 10 tier-A, 313 tier-B, 1,950 tier-C and 17,727 tier-D candidates.

The same framework applies to non-highlighted targets. A user should be able to identify the relevant antigen space, inspect the modeled denominator, recover the target-or peptide-level data, check whether participant-level metadata are public, deidentified or controlled, and trace the displayed value to source data and code provenance. This query path is the practical reason for packaging CHAIR as a Resource rather than as a narrative-only antibody-aging survey.

## Discussion

CHAIR adds an antigen-resolved humoral-memory layer to healthy adult aging resources. Its primary contribution is not a single biomarker, diagnostic model or mechanistic claim, but an integrated Resource framework that links self-antigen, virome and toxin or virulence-factor PhIP-seq with routine clinical screening and T/B/NK immune-cell phenotyping in the same clinically screened cohort. The Resource is intentionally organized around linked matrices, denominator dictionaries, source-data records and access tiers because antibody-recognition histories are only partly captured by host-state measurements made at one sampling time. By pairing antigen-resolved reactivity matrices with cohort metadata, antigen spaces, source-data records, and model outputs, CHAIR makes focal immune-aging features inspectable and reusable rather than leaving them as isolated figure observations.

The central biological pattern is broad antibody-reactome stability with focal age- and sex-aware remodeling. Across the modeled self-antigen, virome and virulence-factor spaces, FDR-supported target-level associations occupied small fractions of the tested antigen universes rather than showing global humoral collapse or uniform expansion. This framing is important for healthy-cohort interpretation. It avoids treating adult aging as a generalized increase in autoreactivity, and it also avoids dismissing sparse target-level signals as noise simply because global summaries remain broadly preserved. Instead, CHAIR supports a reference baseline in which selected self, viral and toxin or virulence-factor entries capture focal shifts in reactivity magnitude, breadth or immunodominance.

This pattern also clarifies how CHAIR should be compared with existing aging resources. Host-state atlases are well suited for measuring contemporaneous cellular, proteomic or clinical states, whereas antibody-reactome profiling adds a memory-weighted antigen-recognition layer. The two views are not interchangeable. By linking antibody reactivity to T/B/NK and clinical measurements in the same cohort, CHAIR allows focal antigen associations to be interpreted against a broader healthy-adult immune context rather than in isolation.

The focused analyses illustrate the balance between utility and boundary. Herpesvirus summaries, including HCMV strata and the VZV query example, are biologically plausible because herpesviruses establish long-lived antigenic memory and population antibody signatures can vary with age, inflammation and host factors^19,20^. In CHAIR, however, these signals remain seroreactivity modules; infection status, viral load, recent exposure, reactivation and menopause-linked mechanisms require orthogonal evidence. The DAZ module also has a plausible biological context because DAZ-family proteins are germ-cell-associated RNA-binding proteins with repeat-rich paralogue architecture^26^. Its male-biased repeat-reactivity pattern therefore merits follow-up in reproductive immunology, but infertility biomarkers, male-exclusive diagnostics, causal reproductive autoimmunity and blood-testis-barrier mechanisms require functional validation. The mimotope workflow prioritizes self-foreign bridge candidates using sequence and aggregate co-reactivity evidence; molecular mimicry, infection-induced autoimmunity and pathogen exposure remain hypotheses for follow-up rather than conclusions from the present data. Similarly, the IAI connects antibody-reactome features with cellular and clinical measurements in an internally cross-validated model, complementing immune-aging clock efforts^6^, but it remains cohort-specific and should not be treated as an externally validated biological-age clock, clinical diagnostic, clinical biomarker or generalizable predictor.

Several limitations define the correct use of CHAIR. First, the cohort is cross-sectional, so within-person aging trajectories, temporal ordering and causal relationships cannot be inferred, and longitudinal validation is still required. Second, CHAIR currently represents a single clinically screened healthy Chinese cohort, and generalizability to other populations, geographic settings, disease states and longitudinal trajectories requires independent testing. Third, PhIP-seq primarily measures antibody reactivity to displayed linear peptides and may miss conformational, glycosylated, lipid-associated or post-translationally modified epitopes. Fourth, target-level attribution can be complicated by homologous proteins, related viral species, shared repeat structures and annotation granularity. Fifth, detailed lifetime exposure histories, latent-infection status, reproductive history, medication use beyond screening exclusions and future health outcomes are not comprehensively resolved, although recent vaccination and recorded infection were excluded during cohort screening. Finally, we do not provide functional or orthogonal binding proof for the focal DAZ or mimotope mechanisms.

These limitations also define the most useful next applications. CHAIR can serve as a healthy-baseline comparator for disease cohorts, vaccine-response studies, infection-history analyses, reproductive immunology studies and population comparisons. It can also support targeted follow-up of focal age-associated self-antigens, herpesvirus seroreactivity modules, DAZ-repeat reactivity, toxin or virulence-factor interactions, high-priority mimotope bridge candidates and multimodal immune-age features. The Resource package organizes processed matrices, metadata, library annotation, target- and peptide-level model outputs, source-data records, data dictionaries, access manifests and figure-code provenance into staged public-summary, deidentified and controlled-access layers. The main claim is therefore deliberately bounded: CHAIR maps focal antibody-reactome remodeling across healthy adult aging and provides a reusable framework for studying antigen-specific immune aging in human populations.

## Methods

### Study design, ethics and participant screening

The Chinese Healthy Aging Immune Resource (CHAIR) was designed as a clinically screened, cross-sectional healthy-adult immune-aging resource. Participants were adults undergoing routine health examinations at the Physical Examination Center of Shanghai Tong Ren Hospital, Shanghai, between January and May 2024. The study was conducted under local institutional ethics approval from Shanghai Tong Ren Hospital (K2024-010-01) and Shanghai Jiao Tong University (B20230280P), and informed consent was obtained before sample collection. The analysis framework was intended to define a clinically screened healthy-adult baseline rather than a disease cohort or clinical diagnostic study.

Clinical screening followed the health-examination and source-protocol criteria used for the CHAIR cohort. Participants with recorded cardiovascular disease, hyperlipidemia, acute or chronic infectious disease, autoimmune disorder or malignancy were excluded according to the source protocol. Participants also had no recorded recent infection or recent vaccination at the time of sampling, reducing the likelihood that small seroreactivity differences reflected acute immune stimulation rather than baseline variation in the healthy-aging cohort. The clinical screening layer included routine health-examination variables spanning complete blood count, urinalysis, liver function, renal function, fasting blood glucose and lipid profiles. These clinical variables were used to support cohort screening, describe the healthy-adult reference range and provide one modality of the multimodal CHAIR data structure; they were not used to assign disease diagnoses in the present manuscript.

### Sample collection and clinical/T/B/NK data capture

Fasting venous blood was collected in the morning as part of the health-examination workflow. Serum-separation tubes (Guangzhou Yangpu Medical Devices Co, cat. no. 623050202) were used for PhIP-seq assays; blood was allowed to clot, centrifuged to isolate serum and stored as serum aliquots at −80°C until analysis. EDTA anticoagulant tubes (Guangzhou Yangpu Medical Devices Co, cat. no. 723050202) were used for complete blood count and flow-cytometry immunophenotyping. Whole-blood T/B/NK phenotyping was performed using antibodies against CD3 (FITC; BD, cat. no. 349201), CD4 (PE-Cy7; BD Pharmingen, cat. no. 557852), CD8 (APC-Cy7; BD Pharmingen, cat. no. 557834), CD19 (APC; BD, cat. no. 340722), CD45 (PerCP-Cy5.5; BD Pharmingen, cat. no. 564105), CD16 (PE; BD, cat. no. 340705) and CD56 (PE; BD, cat. no. 340724).

Samples were acquired using a BD FACSCanto II flow cytometer (Becton, Dickinson and Company) and analyzed using FlowJo software (Waters Biosciences). Whole-blood T/B/NK phenotyping quantified CD3+ T cells, CD3+CD4+ T cells, CD3+CD8+ T cells, CD19+ B cells and CD16+CD56+ natural killer cells as percentage and absolute-count measures. Absolute subset counts were derived from cytometry percentages and complete-blood-count lymphocyte counts. Detailed staining, acquisition, gating and variable-unit information are provided in the clinical and T/B/NK variable dictionary.

### Sample identity harmonization and participant bridge

Sample identities were harmonized before analysis to preserve consistent analysis denominators. The bridge workflow loaded the PhIP-seq, clinical and T/B/NK source datasets; standardized participant identifiers; removed whitespace; removed PhIP-seq technical replicate samples marked by replicate suffixes; collapsed repeated T/B/NK measurements to one participant per physical-examination identifier; checked leading-zero barcode mismatches; added age and sex from the source tables; and classified participants into modality-overlap categories. The workflow exported an audit workbook, validation report, run log and participant bridge used for Supplementary Fig. 1 and Supplementary Table 1.

The denominator-validated participant bridge spans 1,205 unique individuals across available study modalities. The PhIP-seq input contained 1,044 raw records that resolved to 1,008 unique participants after removal of 36 technical replicates. The T/B/NK input contained 999 raw records including 22 replicates that resolved to 977 unique participants. The clinical layer contained 995 unique records without replicate entries in the validated bridge. The principal PhIP-seq analysis base consisted of 1,008 adults aged 20-94 years, including 545 female and 463 male participants. Among the three modalities, 853 participants had PhIP-seq, clinical and T/B/NK data; modality-specific subsets included 153 PhIP-seq-only participants, 55 T/B/NK-only participants, 75 clinical-only participants, 67 participants with clinical and T/B/NK data only and 2 participants with PhIP-seq and T/B/NK data only. No age or sex values were missing in the validated participant bridge.

Because barcode formatting can create false sample mismatches, leading-zero discrepancies were audited separately. Seven T/B/NK barcode records were corrected by adding the leading zero required to match the corresponding PhIP-seq identifier. All seven corrections matched the corresponding PhIP-seq participant by age and sex and were accepted into the corrected bridge. The corrected bridge was then used as the denominator dictionary for participant counts, modality overlap, figure-source data and resource metadata.

### Antigen-library design and count levels

CHAIR used three PhIP-seq antigen libraries to profile antibody reactivity across complementary antigen spaces^14–16,27^: a human self-antigen library, a human virome library and a toxin/virulence-factor library. The human self-antigen library was generated by downloading all available human protein and isoform sequences were downloaded from the NCBI protein database and tiled into 56-amino-acid peptides with at least 28 amino acids of overlap, with increased tiling density at protein termini and exon junctions. The virome library used the same peptide-display strategy for viral and infectious-memory antigens. The toxin/virulence-factor library was constructed from toxin-associated and virulence-factor-associated protein sources curated with toxin/virulence-factor resources and prior environmental-toxin PhIP-seq work^28,29^. All libraries were cloned and packaged into T7 bacteriophage using the T7Select® 10-3 Cloning Kit (Merck), in accordance with the manufacturer’s instructions.

We distinguish raw library and matrix counts from modeled target denominators. The source matrix used for the human self-antigen library contains 646,540 peptides and 646,540 unique peptide identifiers, with 23,710 source-gene labels in the source annotation summary. The virome source matrix contains 122,601 peptides and 122,601 unique peptide identifiers, with 321 source species groups and 1,224 organism or source entries in the library-count summary. The toxin/virulence-factor source matrix contains 173,395 peptides, 172,457 unique peptide identifiers, 1,262 raw target-name labels and 1,482 representative organisms. These source-library counts are distinct from the modeled denominators used for target-level analyses: 23,584 self-antigen targets, 210 virome targets and 1,258 modeled virulence-factor target entries.

### PhIP-seq immunoprecipitation and sequencing

PhIP-seq immunoprecipitation and sequencing were performed separately for each antigen library. Deep-well 96-well plates (Nunc cat. no.278752) were blocked overnight at 4°C with 3% BSA (cat. no. 36101ES80) in PBS, using 800 µL per well, and stored at −80°C until use. Phage libraries were diluted in 0.2% IgG-free BSA (cat. no. 36103ES25) in PBS to approximately 16,000-fold library diversity per milliliter. For each selection, 10 µL serum, corresponding to approximately 2 µg IgG, was added to the diluted phage library and incubated for 16 h at 4°C with gentle end-over-end mixing. Buffer-equilibrated rProtein G MagPoly beads (Smart-Lifesciences, cat. no. SM036050; 40 µL per well) were then added, and samples were incubated for 4 h at 4°C with rotation.

After immunoprecipitation, unbound phage was removed and beads were washed four times with 200 µL TBS containing 0.1% IGEPAL CA-630 (Sigma-Aldrich, cat. no. I3021-100ML). After the first wash, beads were transferred to PCR plates. After the final wash, beads were resuspended in 40 µL sterile water and transferred to a new 96-well PCR plate. Phage particles were heat-treated at 95°C for 10 min to release DNA and stored at −80°C for up to one week or processed directly for library preparation. The first PCR used Q5 High-Fidelity DNA Polymerase (New England Biolabs, cat. no. M0491L) for 28 cycles to introduce sample-specific barcodes. The second PCR used Q5 polymerase for 8 cycles to introduce plate-specific indexes. PCR success was assessed by agarose gel electrophoresis on selected reactions. Pooled PCR products were resolved on a 2% agarose TAE gel, recovered using a gel extraction kit (FlaPure Gel Purification Kit, Gensand, cat. no. GE706), and, after concentration measurement, pooled for next-generation sequencing on the Illumina novaseq Xpuls platform.

### Read processing, normalization and enrichment-factor scoring

Raw sequencing reads were demultiplexed by barcode indexes and mapped to the corresponding peptide libraries. Raw read counts were normalized within each sample by dividing each peptide count by the total peptide count for that sample and scaling to 1.25 million reads. For each plate, five bead-only control selections were included.

Normalized bead-only controls were compared by Pearson correlation, and at least three controls with r > 0.5 were required to calculate the mean bead-only baseline for a plate. Peptide enrichment factors (EFs) were calculated by dividing the normalized peptide count in each sample by the mean normalized bead-only count for the same peptide (library and selection QC are described in supplementary Figure S2).

Library-specific enrichment thresholds were determined using kernel density estimation and Gaussian mixture modeling to balance likely antibody-enriched peptides against background signal. For each sample and library, enriched-peptide fractions, enrichment-factor summaries and library-appropriate breadth or intensity metrics were calculated. Human self-antigen peptides were summarized to gene or modeled target labels. Virome peptides were summarized to species or viral target labels with common-name annotations, and toxin/virulence-factor peptides were summarized to target names and representative organism or source-class labels. Closely related strains or annotation entries were collapsed to higher-level viral, organism or target labels where specified in the source tables.

### Target-level aggregation across antigen libraries

Processed PhIP-seq enrichment-factor matrices were converted into sample-level target summaries before cross-library modeling. The aggregation rules were defined separately for each antigen library because the self-antigen, virome and toxin/virulence-factor libraries differ in peptide design, annotation structure and target granularity. The human self-antigen library was summarized to gene or protein target labels using greedy Top10 peptide panels generated within each target. Peptides were initially ranked by a candidate score equal to positive-sample fraction multiplied by ln(1 + summed EF among positive samples). Within up to 300 highest-ranked individual candidates per target, a greedy algorithm iteratively selected the peptide adding the largest number of previously uncovered positive samples, using candidate score to break ties; selection stopped after 10 peptides or when no additional sample coverage was gained. For each modeled self-antigen target, the primary target score was the mean log1p-transformed EF across the selected Top10 panel, and target positivity was defined by at least one panel peptide with EF > 3.14. The modeled self-antigen universe contained 23,584 targets. This EF > 3.14 cutoff applies to target-panel construction and target positivity; the peptide-level regression described below used EF > 4.93 for the human self-antigen library.

The virome and toxin/virulence-factor libraries were summarized using target panels designed to capture recognition of a viral species, viral target label, source target or representative organism/source-class entry. For the virome library, candidate panels were generated from up to 300 individually ranked candidates per target using a specificity-weighted candidate score and greedy incremental sample coverage; the first five selected peptides formed the Top5 target panel. Each virome target-level score was the maximum log1p-transformed EF across the Top5 panel, and target positivity was defined by at least one Top5 panel peptide with EF > 3.14. The modeled virome universe contained 210 target/species-group units. For the toxin/virulence-factor library, up to 350 ranked candidates per target were considered, and a greedy selector combined incremental sample coverage with specificity-tier and protein-diversity terms to construct panels of up to 10 peptides; the first five formed the Top5 score panel. Each toxin/virulence-factor target-level score was the maximum log1p-transformed EF across that Top5 panel, and target positivity in the harmonized sample-level matrix was defined by at least one Top5 panel peptide with EF > 3.74. The modeled toxin/virulence-factor universe contained 1,258 target entries. For source traceability, the descriptive positive_fraction field exported for toxin/virulence-factor rows in Supplementary Table 3 retains the upstream Top10 panel coverage; it is distinct from the Top5 positivity matrix used for target-score modeling and interaction reruns. These aggregation choices retain library-specific biological and technical structure while producing harmonized target-level fields for downstream age, sex and interaction analyses.

### Global antibody-reactome summaries

Global antibody-reactome summaries were calculated within each antigen domain using the 1,008-sample PhIP-seq analysis base. Age-group and sex summaries were derived from the harmonized sample metadata and target-level matrices. Because the three libraries differ in peptide density and target definition, global summaries were interpreted within antigen domain rather than as a single shared breadth scale. Self-antigen summaries used sample-level and target-level breadth or score distributions. Virome and toxin/virulence-factor summaries used target-level positive fractions, score distributions and positive-peptide or positive-target summaries as appropriate for the figure-source layer.

Global-to-focal summaries combined library-wide target denominators with the number and fraction of targets passing false-discovery thresholds for age, sex or age-by-sex terms. These summaries were used to describe whether age-, sex-or interaction-associated patterns were broad across a library or concentrated in a small focal tail of targets. The manuscript reports focal counts and fractions by library, preserving the target denominators of 23,584 self-antigen targets, 210 virome targets and 1,258 toxin/virulence-factor targets.

### Target-level age, sex and interaction models

Harmonized target-level model outputs were generated for the 1,008-sample PhIP-seq base set. For each modeled target, the analysis table records the antigen library, target identifier, target name, common-name annotation where available, model level, sample count, median logEF score, upper-tail score summary, positive fraction, peptide-panel size or candidate count, age effect, sex effect, age-by-sex interaction fields where available, P values, FDR-adjusted q values and source-file provenance. Because the three target-level source analyses used library-specific models, inferential procedures were not identical across antigen spaces. For human self-antigen targets, age effects were sex-adjusted partial Pearson correlations between chronological age and the Top10 mean log1p(EF) score; sex effects were male-minus-female coefficients from ordinary least squares (OLS) models adjusted for standardized age; and age-by-sex effects were interaction coefficients from OLS models containing standardized age, sex and an age-by-sex term. Human target-level coefficient P values were evaluated with t distributions using residual degrees of freedom. For virome and toxin/virulence-factor Top5 panels, primary age effects were Spearman correlations between target score and age, whereas sex effects were estimated by OLS models of standardized target score containing standardized age, age squared and sex. Virome and toxin/virulence-factor age-by-sex reruns used OLS models of standardized target score on standardized age, sex and their interaction; secondary linear-probability models evaluated panel positivity. These reruns used two-sided normal-approximation P values.

False-discovery control was applied separately within each antigen library and model term using the Benjamini-Hochberg procedure^29^, and the resulting q values were recorded in the harmonized model table. Unless otherwise stated, targets flagged at the FDR 0.05 level were treated as statistically supported target-level associations. Age, sex and age-by-sex terms were interpreted separately, and an “any focal FDR” summary was used only to describe the union of targets with at least one FDR-supported term. Effect directions were summarized separately for age-increasing and age-decreasing targets. The source package records the target-level outputs (Supplementary Table 3).

The target-level model layer was designed as an association screen within a clinically screened healthy-adult resource. Target-level virome and toxin/virulence-factor associations were interpreted as antibody-reactivity summaries, not as evidence of active infection, viral load, recent exposure or organism-specific diagnosis. Self-antigen associations were interpreted as population-level seroreactivity features, not as evidence of clinical autoimmunity.

### Peptide-level regression and source trace

Peptide-level regression was used as a source-trace layer for target-level and figure-level claims. Peptides were included in the regression table after library-specific positivity filtering. Supplementary Table 4 records 227,850 tested human self-antigen peptides from 646,540 peptides using an EF positivity cutoff of 4.93 and a minimum-positive filter of 0.05, corresponding to 53 positive samples in the 1,008-sample base. The virome analysis tested 4,484 of 122,601 peptides using an EF positivity cutoff of 3.14 and the same 0.05 minimum-positive filter. The toxin/virulence-factor analysis tested 36,160 of 173,395 peptides using an EF positivity cutoff of 3.74 and the same 0.05 minimum-positive filter. Log_2_(EF + 1)-transformed enrichment factors were modeled using ordinary least squares linear regression with age (mean-centered, continuous) and sex as predictors. Wald χ² tests were used to assess the significance of the age coefficient and the overall effect of sex. P values were adjusted for multiple testing within each library and separately for age and sex using the Benjamini–Hochberg false discovery rate (FDR) correction.

For each tested peptide, regression outputs record the peptide identifier, source annotation, number of samples, number and fraction of positive samples, positivity cutoff, minimum positive count, Log_2_(EF + 1)-transformation, age-centering value, sex reference, age coefficient per year, age Wald chi-square statistic, age P value, Benjamini-Hochberg q value within library^30^, sex Wald chi-square statistic, sex P value, sex q value within library, residual sigma and fitted coefficients. The peptide-level table is used to trace selected target-level signals back to supporting peptide data and to support figure source-data traceability. It is not used to infer active infection, validated exposure history, clinical diagnosis or molecular mechanism without orthogonal validation.

### Structural mapping and solvent-accessibility summaries

Structural mapping was used to provide biophysical context for high-sample-positivity PhIP-seq peptides. Here, high sample positivity denotes peptides selected from the most sample-positive candidates within each antigen library for which a preferred structural mapping could be retained; it is not used as a universal cross-library positivity cutoff.

Candidate peptides were selected from the human self-antigen, virome and toxin/virulence-factor libraries, and one preferred structural mapping was retained for each mapped peptide. Peptide-level surface annotations were derived from AlphaFold Database^31–33^ (accessed between 21 and 24 July 2026), structure mappings with FreeSASA/RSA output^34^, and residue-level annotations were summarized to peptide-level metrics. The primary mapped set contained 412 peptides, comprising 142 human self-antigen peptides, 145 virome peptides and 125 toxin/virulence-factor peptides.

Because some virome peptides lacked direct high-stringency structure matches, the primary analysis included a relaxed virome mapping layer. A strict-only sensitivity set retained original mapping-threshold entries with coverage >= 0.95 and identity >= 0.85, yielding 314 mapped peptides comprising 142 human self-antigen peptides, 47 virome peptides and 125 toxin/virulence-factor peptides. Mapping quality, partial-coordinate status, positive-sample fractions and the 98 relaxed virome mapping entries are recorded in Supplementary Table 5.

For each mapped peptide, surface summaries included mean solvent-accessible surface area, mean relative solvent accessibility (RSA), maximum RSA, the percentage of annotated residues classified as surface exposed, mean model-confidence score where available and largest surface patch metrics^35^. Residues were classified as buried when RSA < 0.05, intermediate when RSA was >= 0.05 and < 0.20, and exposed when RSA >= 0.20^25^. Descriptive summaries reported n, mean, s.d., minimum, quartiles, median, maximum and interquartile range for each library and mapping cohort.

Across-library RSA and surface-summary comparisons used Kruskal-Wallis tests for the three antigen-library groups, with Benjamini-Hochberg FDR correction^30^ across six prespecified outcomes within each mapping cohort. Pairwise two-sided Mann-Whitney U tests with continuity correction were performed only for metrics with omnibus results meeting the prespecified FDR threshold, and Holm correction was applied across triggered pairwise follow-up tests within each cohort. Effect sizes were recorded as epsilon-squared for omnibus tests and rank-biserial correlations for pairwise contrasts.

These structural analyses were interpreted as context for selected mapped peptides, not as evidence of antigen exposure, antibody-binding mechanism or causal biology.

### Structure-matched permutation testing

Structure-matched permutation tests asked whether selected mapped peptides had greater solvent accessibility than expected from same-structure background windows. For each mapped peptide, the null background was generated by drawing uniform contiguous windows within the peptide’s own mapped structure, with window length equal to the aligned annotated residue span. This design preserved the source structure and structural peptide length while randomizing the peptide window location. The permutation analysis evaluated mean peptide-level RSA and mean percentage of surface-exposed residues, defined as residues with RSA >= 0.20.

For each cohort and metric, 10,000 random windows were generated with seed 20260724. One-sided empirical P values tested whether the observed statistic exceeded the structure-matched null distribution and were calculated as (1 + number of null statistics greater than or equal to the observed statistic) divided by (1 + number of permutations). Benjamini-Hochberg FDR^29^ was applied across eight tests within each primary and strict/sensitivity cohort set. The source package records permutation outputs, null summaries and histogram-bin tables (Supplementary Table 5).

### Herpesvirus and HCMV module analyses

Focused herpesvirus analyses summarized seroreactivity patterns using non-saturated metrics rather than binary positivity alone. For each annotated herpesvirus group, sample-level metrics included positive-peptide breadth, summed positive EF, maximum EF, strict positivity indicators and high-response strata based on upper-tail breadth, summed-EF or maximum-EF thresholds. Age-bin, sex and age-bin-by-sex summaries were calculated using the 1,008-sample PhIP-seq analysis base, with stratum-specific sample counts retained in the source tables.

HCMV analyses used the same seroreactivity framework, with additional checks to avoid overinterpreting near-saturated binary positivity. Summary tables recorded any-positive and strict-positive percentages, breadth, protein breadth, summed positive EF, maximum EF and high-response strata. Focused HCMV checks included continuous-age rank correlation, nonlinear age-bin permutation tests, sex-difference permutation tests for breadth and magnitude metrics, and upper-tail high-response risk-ratio or odds-ratio summaries where specified. Midlife female-versus-male contrasts were reported for strata with balanced sample counts and were treated as seroreactivity breadth and magnitude contrasts. HCMV and broader herpesvirus metrics were not interpreted as active infection, viral load, recent exposure, individual infection status, viral reactivation or a menopause mechanism.

### DAZ repeat and virulence-factor interaction modules

DAZ analyses summarized self-antigen reactivity to DAZ-family peptides and repeat-rich DAZ segments. Sample-level DAZ metrics included any DAZ positivity, DAZ positive-peptide count, DAZ summed positive EF, DAZ maximum EF, DAZ-repeat positive-peptide count, DAZ-repeat summed positive EF, DAZ-repeat maximum EF, and upper-quartile or upper-decile high-response flags. Binary DAZ positivity was retained as context because it was broadly prevalent, whereas repeat-focused magnitude metrics were used to describe the focal module. Age-bin, sex and age-bin-by-sex summaries retained stratum-specific sample counts.

DAZ repeat architecture was summarized from peptide metadata mapped to DAZ2, DAZ3 and DAZ4 and to canonical DAZ repeat strings^24^. Peptides were classified as repeat-rich or non-repeat-rich for architecture summaries, and DAZ paralogue homology was treated as a source of shared repeat-sequence interpretation rather than as evidence that a signal uniquely identifies one paralogue. DAZ inferential summaries included sex and age-stratum contrasts, bootstrap confidence intervals for median differences, permutation tests for log1p mean differences and permutation tests for upper-tail response proportions with multiple-testing correction by metric. These analyses were interpreted as repeat-epitope antibody-reactivity patterns in a subset of participants, not as male-exclusive diagnostic markers, infertility biomarkers, causal reproductive autoimmunity, blood-testis-barrier mechanisms or Y-linked disease mechanisms.

Virulence-factor interaction modules were drawn from the harmonized target-level interaction fields and focused source tables. Score-level and positivity-level age-by-sex interaction summaries were kept distinct from main sex-effect summaries. Virome and toxin/virulence-factor interaction checks were interpreted as resource features for follow-up rather than as evidence of sex-specific toxin, pathogen or environmental exposure.

### Mimotope bridge candidate generation

Mimotope bridge analysis was used to nominate cross-antigen peptide pairs linking human self-antigen candidates with virome or toxin/virulence-factor candidates profiled in the same PhIP-seq cohort. The analysis was designed as a prioritization workflow for follow-up hypotheses, not as a validation assay for molecular mimicry, pathogen exposure or infection-induced autoimmunity.

Human query peptides were drawn from the human self-antigen candidate atlas by retaining the top ten candidate peptides per protein with positive-sample fraction >=0.05 in the 1,008-sample PhIP-seq analysis base. Foreign query peptides were drawn from the virome and toxin/virulence-factor candidate atlases by retaining peptides with positive-sample fraction >=0.01 in the same base sample set. Peptide sequences were normalized before matching by removing non-amino-acid characters and obvious synthetic linker runs where present. Low-complexity sequences were flagged using residue-composition and entropy rules, and low-complexity 7-amino-acid seeds were not used as the sequence entry point.

The workflow scanned 160,072 human candidate peptides and indexed 20,295 virome or toxin/virulence-factor candidate peptides. Candidate sequence bridges were generated by searching for shared exact 7-amino-acid seeds, extending each shared seed to its maximal exact matched span and to a biochemically similar span based on amino-acid similarity classes. High-occurrence seed removal and a per-human-peptide cap were used to limit redundant sequence matches; no high-occurrence seeds were removed in this run, and up to eight foreign peptide matches were retained for each human peptide before co-reactivity scoring. This produced 24,501 sequence candidate pairs for co-reactivity evaluation.

### Co-reactivity scoring and homology-risk audit

For each candidate pair, log-transformed enrichment-factor vectors were compared across the 1,008 base samples. Human and virome positive calls used EF > 3.14, and toxin/virulence-factor positive calls used EF > 3.74, matching the library-specific positivity rules used elsewhere in the resource. Co-reactivity features included Pearson correlation of logEF vectors, the number and fraction of samples positive for both peptides, expected positive-overlap count under independence, positive-overlap enrichment, positive-overlap odds ratio and positive-overlap Jaccard index.

Bridge scoring combined sequence similarity and aggregate co-reactivity while penalizing likely background explanations. The sequence component incorporated exact span length, shared-seed multiplicity, human and foreign candidate scores, positive fractions and low-complexity penalties. The final bridge score added non-negative Pearson logEF correlation, enrichment above the independent-overlap expectation, both-positive sample count and additional biochemical similarity beyond the exact span, then subtracted penalties for low-complexity sequence context, long exact-homology windows, nonpathogen metazoan background entries and moderate or high protein-homology risk.

Protein-homology risk was assessed as a local annotation-based audit rather than a full-length homology search. The audit flagged nonpathogen metazoan background entries, exact peptide spans of at least 20 amino acids, extended peptide matches with overlapping protein-function tokens where annotation was available, eukaryotic pathogen homolog backgrounds with extended similarity, and toxin/virulence-factor entries requiring annotation review. These flags were used to down-weight or remove peptide pairs from the primary display layer, not to prove or exclude structural or evolutionary homology.

### Evidence tiers and hardened bridge tables

Candidate pairs were ranked by evidence priority and bridge score, and the final reported bridge table was limited to 20,000 pairs. Evidence tiers were assigned before secondary review. Long exact-homology windows and nonpathogen metazoan background entries were assigned to audit/background categories outside the main A/B evidence classes before A/B tiering. Tier A required no low-complexity warning, no long-homology or audit-background flag, exact span >=8 amino acids, biochemical span >=10 amino acids,

Pearson logEF >=0.15, positive-overlap enrichment >=1.25 and both-positive count >=20. Tier B used the same audit prefilter and required no low-complexity warning, exact span >=8 amino acids, both-positive count >=10, and either Pearson logEF >=0.10 or positive-overlap enrichment >=1.20. Tier C retained non-low-complexity sequence-only follow-up candidates with exact span >=9 amino acids and limited co-reactivity support. Tier D contained lower-priority low-complexity or weak co-reactivity.

The reported table contained 10 tier-A pairs, 313 tier-B pairs, 1,950 tier-C pairs and 17,727 tier-D pairs. A/B evidence therefore comprised 323 candidate pairs. A secondary-review layer retained A/B candidates with co-reactivity support and without low-complexity or moderate/high protein-homology-risk flags, producing 316 candidates. A stricter primary-keep layer then removed long exact/biochemical homology and repeat-like shared-motif flags, yielding 251 pairs for focused display and resource navigation.

The source package records the full candidate table, filter funnel, evidence-tier definitions, collapsed human-target and foreign-target summaries, motif summaries, network summaries and claim-boundary fields (Supplementary Table 7).

All mimotope bridge outputs were interpreted as sequence-similarity and cohort-level co-reactivity prioritization evidence. They do not establish antigenic cross-reactivity, antibody-clone sharing, active or historical infection, pathogen exposure, clinical autoimmunity, diagnosis, treatment targets or causal immune mechanisms without orthogonal validation.

### Immune Age Index modeling overview

Immune Age Index (IAI) analyses were designed as an internal resource use case for summarizing multimodal immune-aging structure in CHAIR. Chronological age was used as the modeling outcome. Candidate predictors came from clinical and lymphocyte measurements, human self-antigen PhIP-seq features, virome PhIP-seq features and toxin or virulence-factor PhIP-seq features. Model performance was summarized by mean absolute error (MAE), root mean squared error (RMSE), Pearson correlation between predicted and chronological age, fold-local mean-age baseline MAE and MAE improvement relative to that baseline. Preprocessing and model fitting were fold-local; feature selection was fold-local where specified below, whereas fixed-panel and fixed-pool analyses used preselected feature sets. The IAI was interpreted as cohort-specific internal validation rather than as an externally validated biological-age clock.

### Compact complete-case model

The compact IAI model used a complete-case modeling framework to retain interpretability of a small multimodal panel. Candidate features were first ranked within each data layer by the absolute strength of their sex-adjusted association with chronological age. Peptide features were represented as log-transformed enrichment-factor measurements, and clinical or lymphocyte features were taken from the harmonized clinical/T/B/NK feature maps. Estimated GFR CKD-EPI and estimated GFR MDRD were excluded before feature ranking because they are derived renal-function estimates with strong age dependence. The top 100 available features per layer were retained for compact-panel search, yielding 345 candidate features across the available layers.

Compact feature combinations were explored in a complete-case framework requiring at least 100 participants for each evaluated feature set. Five outer folds were generated by stratifying participants by sex and age quantile. Within each training fold, feature preprocessing included fold-local handling of missingness, centering and scaling, and orientation of features according to their training-fold association with sex-residualized chronological age. A beam-search strategy evaluated candidate equal-weight scores by training-fold calibration to chronological age and held-out fold prediction. The resulting compact panel contained 18 features and 115 complete-case participants and was then fixed for the regression-model comparison.

The fixed 18-feature panel was evaluated by regression-model comparison across penalized or linear, kernel, boosting and tree-based regressors. For the regression comparison, preprocessing was performed within each training fold using median imputation and feature standardization followed by model fitting. Hyperparameters were selected inside training folds using inner cross-validation with MAE as the optimization criterion. The final compact IAI model used ElasticNet regularization^36^, building on lasso shrinkage^37^ and implemented through scikit-learn^38^, because it had the lowest cross-validated error in the fixed-panel comparison. In the selected ElasticNet model, the fold-level best hyperparameters were alpha = 0.1 and l1 ratio = 0.1 in all five folds. A parallel sensitivity model added sex as an additional covariate to the same 18 features.

### Near-full nested cross-validation and internal holdout

To test whether broader age-associated structure was present beyond the complete-case subset, we implemented a near-full internal validation workflow across the 1,008-sample PhIP-seq analysis base. Five folds were generated using age-quantile and sex-stratified assignment with random seed. Within each outer training fold, feature screening was performed de novo using only training samples. Clinical and T/B/NK measurements were not required to be complete for all 1,008 participants. Within each outer training fold, each clinical or T/B/NK candidate was screened using the training samples with non-missing values for that feature, with a minimum of 50 available training samples. The screening step selected the top 30 human self-antigen PhIP-seq peptides, top 30 virome PhIP-seq peptides, top 30 toxin or virulence-factor PhIP-seq peptides and top 20 clinical or T/B/NK features by absolute sex-residualized correlation with chronological age. This yielded 110 candidate features per fold.

After feature screening, near-full models used a pipeline consisting of median imputation, standardization and ElasticNet regression33-35; imputation and standardization were fitted within the training data and then applied to the corresponding held-out samples. ElasticNet hyperparameters were tuned inside each outer training fold by grid search over alpha values 0.01, 0.1, 1.0 and 10.0 and l1 ratios 0.1, 0.5 and 0.9, using shuffled 3-fold inner cross-validation and negative MAE as the scoring function. The primary near-full nested-CV summary was calculated from out-of-fold predictions across all five outer folds. Baseline predictions for each test fold were the mean chronological age in the corresponding training fold. The source package records fold metrics, out-of-fold predictions and selected-feature audit rows (Supplementary Table 8).

A separate near-full internal holdout layer used one age- and sex-stratified outer fold as the holdout set. Feature selection, imputation, standardization and ElasticNet tuning were performed using only the remaining training samples, with 4-fold inner cross-validation for tuning, and the fitted model was then evaluated in the held-out samples. This layer used 802 training samples and 206 holdout samples and was treated as an internal holdout from the same cohort, not as an independent external validation cohort.

### Fixed-pool permutation control

The near-full workflow also included a fixed selected-feature-pool age-label permutation control. The feature matrix was held fixed to the holdout-selected feature pool, and chronological-age labels were permuted within sex strata for 100 permutations. For each permutation, models were refit across the same five outer folds using the fixed feature matrix, median imputation, standardization and ElasticNet tuning. This control tested model calibration after selected feature-pool construction; it did not rerun fold-local feature screening and therefore was not a full re-screening permutation pipeline.

Selected features from the near-full workflow were treated as fold-local candidates, not as a locked stable 110-feature signature. Antigen features in compact or near-full IAI models were interpreted as age-informative model features or candidate immune-aging correlates, not as causal antigen exposures, active or historical infection markers, disease mechanisms, clinical diagnostic markers, clinical biomarkers, treatment targets or externally generalizable predictors. The compact model, near-full nested-CV workflow, internal holdout, selected-feature audit, prediction tables and permutation controls are recorded in Supplementary Table 8.

### Statistical analysis and reproducibility

Statistical analyses were performed using R 4.4.3 and Python 3.12 with the software packages specified in the Code availability statement. Unless otherwise stated, statistical tests were two-sided and were evaluated at an alpha level of 0.05. False-discovery rate or family-wise error correction was applied as specified for each analysis. Sample sizes (n) refer to independent participants or independent peptide or target units, as specified for each analysis; technical replicates were excluded from the primary participant-level analyses. Descriptive statistics report the relevant sample size, measure of central tendency and measure of variability, with error-bar definitions provided in the corresponding figure legends. Statistical assumptions and the rationale for parametric or non-parametric tests are described within the relevant analysis subsections. Empirical permutation tests were one-sided where explicitly stated and used prespecified random seeds and permutation counts.

## Data availability

A versioned CHAIR v1.0 original data, Library metadata, Processed PhIP-seq enrichment matrices, sample-level metadata, clinical variables, T/B/NK lymphocyte measurements and Immune Age Index predictions contain participant-level immune, demographic or clinical information and are released only as approved deidentified files or through controlled access, as specified in the final access-tier manifest. Access to controlled participant-level or identifier-linked data requires a scientifically justified request to Sheng-Ce Tao, institutional approval where applicable and a data-use agreement consistent with the study consent and ethics approvals.

## Code availability

All computational analyses were performed using Python 3.12 and R 4.4.3. Python analyses used NumPy v1.26.4, pandas v2.2.2, SciPy v1.13.0, statsmodels v0.14.2, scikit-learn v1.7.1, scikit-posthocs v0.11.2, Matplotlib v3.8.4, Seaborn v0.13.2, Plotly v6.6.0 and Streamlit v1.55.0. Reproducibility code for figure source-data generation is available upon request.

## Resource availability

CHAIR v1.0 is the manuscript-associated resource release for this study. The release is organized around Supplementary Tables 1-8, comprising resource manifests, data dictionaries, access tiers, processed data layers and analysis outputs. The authoritative versioned release is hosted at http://47.239.208.106/.

## Author contributions

G.N.S., H.S. and S.C.T. conceptualized the study. M.C., J.Y., X.S. and H.S. collected cohort samples. J.D. constructed the human proteome library. G.N.S., H.X., J.Y. and X.S. performed experiments. H.Q., M.M., Y.L. and S.G. provided experimental expertise. G.N.S., J.W., M.C., J.Y., J.L., C.H. and Z.S. analyzed the data. Z.S. developed the CHAIR resource infrastructure. H.S. and S.C.T. provided funding and supervision. G.N.S. wrote the initial draft with contributions from J.Y., J.L., Z.S. and C.H. G.N.S., M.C., J.Y., J.L., C.H., Z.S., H.S. and S.C.T. revised the manuscript.

## Acknowledgements

This work was supported by the National Natural Science Foundation of China (Nos. 92374110, 82570838 and 82070730), the National Key Research and Development Program of China (Nos. 2023YFC2307200, 2024YFA1307600, 2024YFA1307601, 2024YFA1307602, 2024YFA1307603 and 2024YFA1307604), the Prevention and Control of Emerging and Major Infectious Diseases-National Science and Technology Major Project (No. 2025ZD01999900), the Major Project of Guangzhou National Laboratory (No. GZNL2023A01005) and the Fundamental Research Funds for the Central Universities (No. YG2023ZD04). The funders had no role in study design, data collection, analysis and interpretation of data, or writing of the manuscript.

