## Supplementary information Figure S1-12 for "CHAIR an antigen-resolved antibody-reactome resource for healthy adult immune aging"

### Content:

Supplementary information Figure S1-12

Supplementary tables 1-10 are provided as separate excel workbooks

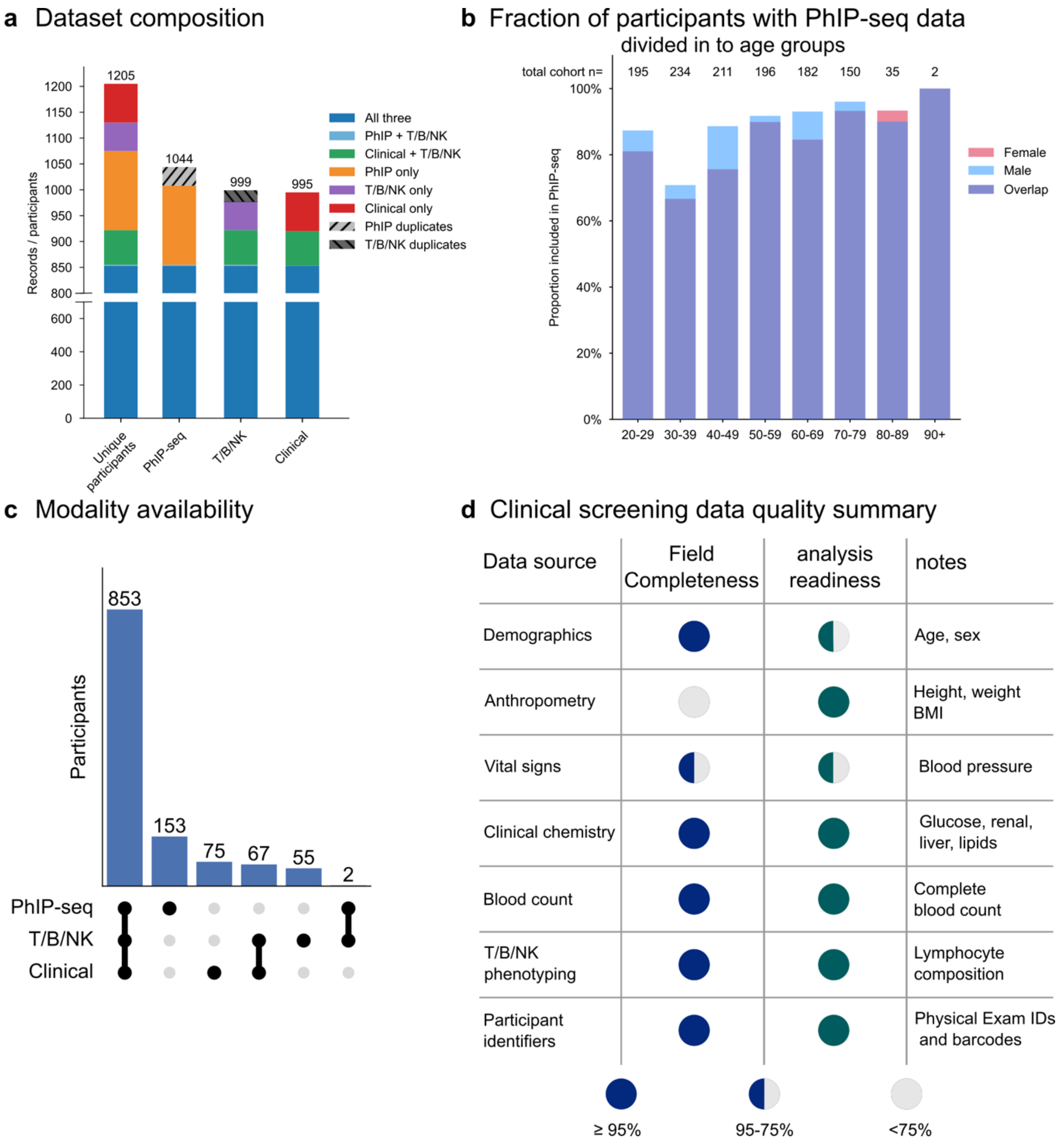


Figure S1. Composition, overlap, and demographic representation of the study cohort across PhIP-seq, T/B/NK immunophenotyping, and clinical screening datasets.

a) Overview of participant composition across the three primary study datasets. The integrated cohort comprised 1,205 unique participants, of whom 853 had data available in all three modalities (PhIP-seq, T/B/NK immunophenotyping, and clinical screening). Individual source datasets contained 1,044 PhIP-seq records representing 1,008 unique participants (36 replicates), 999 T/B/NK records representing 977 unique participants (22 replicates), and 995 clinical screening participants without duplicate entries. Colored segments indicate the contribution of each participant subset to the total cohort, while hatched segments denote duplicate sample records within the PhIP-seq and T/B/NK datasets that were excluded from participant-level analyses. b) Proportion of participants with available PhIP-seq data stratified by age group and sex. Numbers above each bar indicate the total number of participants within each age category. Shaded regions represent the fraction of participants in each demographic stratum with PhIP-seq measurements, illustrating consistently high PhIP-seq coverage across most age groups. The overlap component denotes participants of each sex with available PhIP-seq data, whereas the upper colored segments indicate participants lacking PhIP-seq data within each demographic category. c) Participant overlap between data modalities. UpSet plot summarizing the intersection of participants across PhIP-seq, T/B/NK immunophenotyping, and clinical screening datasets. The largest intersection comprised 853 participants with complete multimodal data. Modality-specific subsets included 153 participants with PhIP-seq only, 75 with clinical screening only, 55 with T/B/NK immunophenotyping only, 67 with both clinical screening and T/B/NK immunophenotyping but no PhIP-seq, and 2 participants with both PhIP-seq and T/B/NK immunophenotyping but without clinical screening. No PhIP-seq + clinical-only participants were identified in the current T/B/NK-based bridge. d) Summary of completeness and analysis readiness for clinical screening variables used in this study. Data domains are grouped by source and evaluated according to field completeness and suitability for downstream analyses. Demographic variables (age and sex), clinical chemistry, complete blood count, T/B/NK immunophenotyping, and participant identifiers were essentially complete, whereas anthropometric measurements and vital signs exhibited lower completeness but remained suitable for analysis after quality control. Filled circles indicate high completeness or readiness, partially filled circles indicate intermediate completeness, and open circles indicate lower completeness, as summarized in the legend.

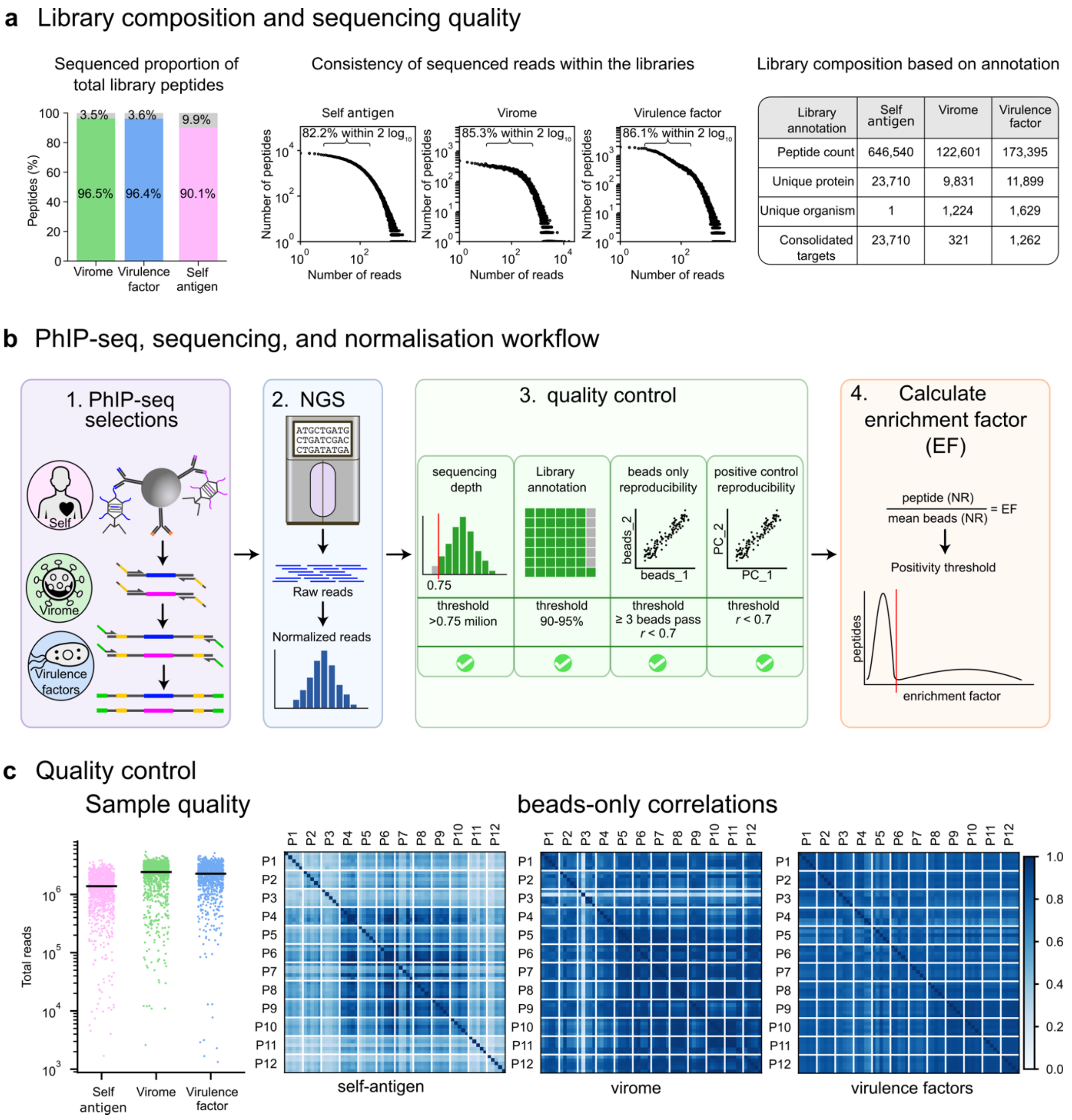


### Figure S2. PhIP-seq quality control

a) Three libraries used in selection. The Human self-antigen library contains 646,540 peptides, 90.1% of peptides have been verified through library sequencing and 82.2% of peptides are sequenced within ± 1 log. The Virome library contains 122,601 peptides and 96.5% of them are verified through sequencing. The Virulence factor library contains 173,395 peptides and 96.4% have been verified through sequencing. b) PhIP seq, sequence analysis, and QC pipeline. c) The median sequencing depth of each sample is above 1 million in all libraries. The Beads only correlation quality control shows that the smaller libraries have better internal correlation than the larger Human self-antigen library.

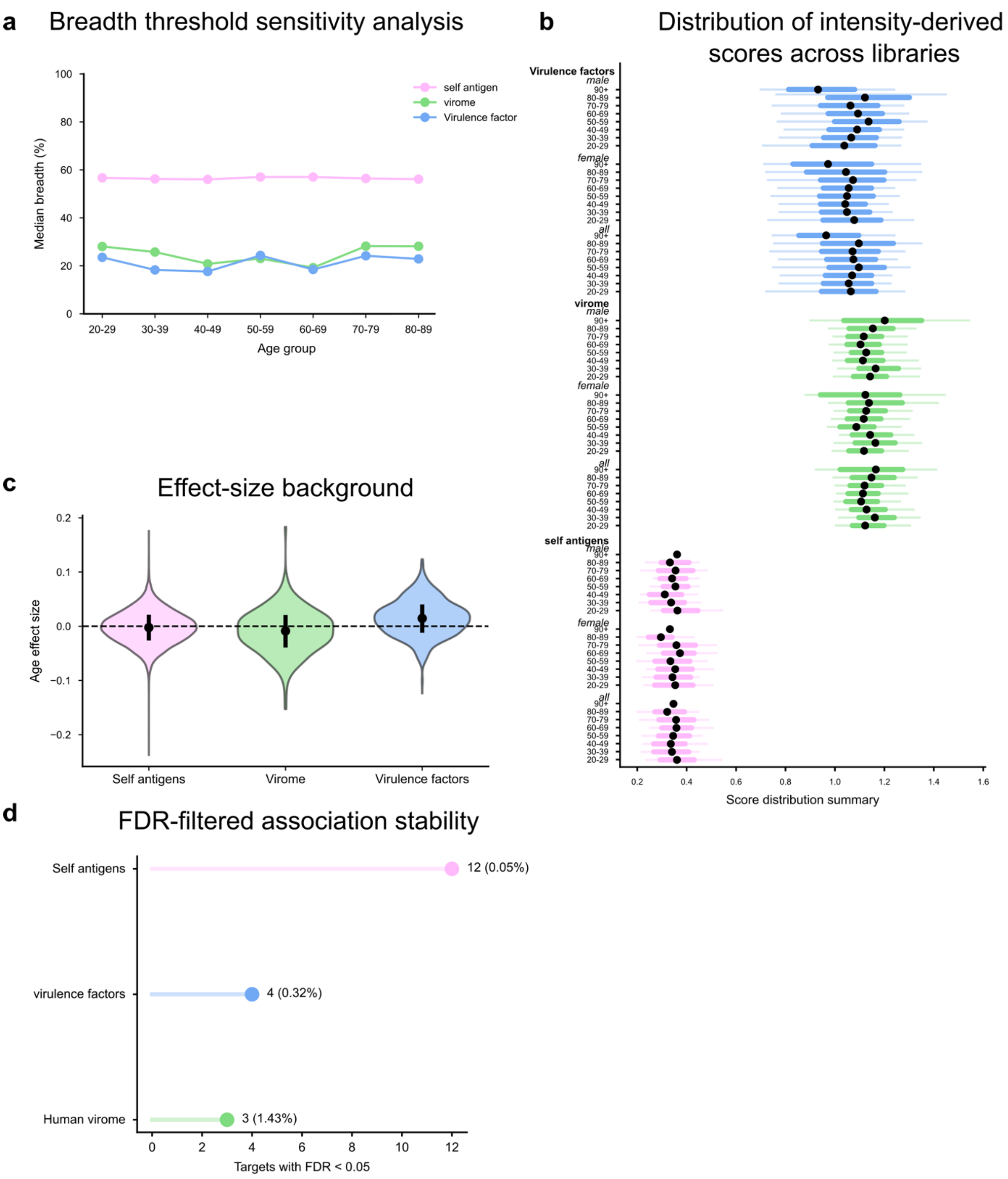


Figure S3. PhIP-seq selections summary

a) The median breadth of each library is stable across age groups but generally higher in the Self-antigen library. b) Distribution of intensity derived scores per library. Human library uses mean EF as score and VF and virome library uses max EF as metric. Generally the strength is stable across age groups. c) Distribution of background age effect sizes across antigenic libraries. Violin plots depict the density of peptide-level age effect estimates across the self-proteome, virome, and toxin/virulence factor antigen spaces. Median values are shown as black points with interquartile ranges, with the dashed line indicating an effect size of zero. d) Stability of antigen associations after FDR correction. Horizontal dot plots summarize the number of targets with significant associations (FDR < 0.05) identified within the self-antigen, human virome, and virulence factor libraries. Percentages indicate the fraction of total targets within each antigen space passing the significance threshold.

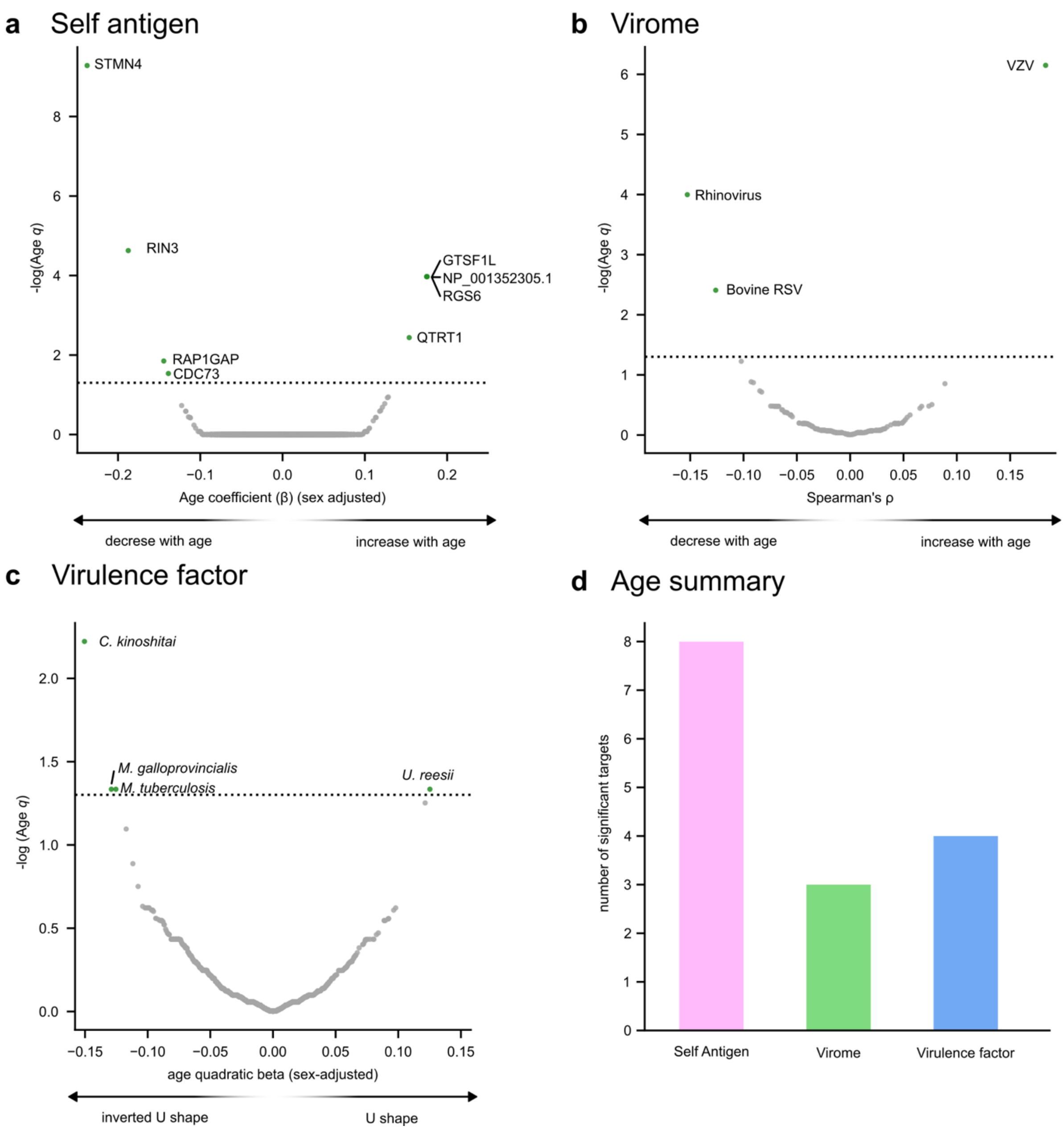


Figure S4. Age-associated antigenic signatures across self-antigen, virome, and virulence factor spaces.

a)–c) Volcano plots showing age-associated associations across three antigen libraries. Each point represents an individual antigen target, with the x-axis showing the age association metric: sex-adjusted age regression coefficient (β) for self-antigens a), Spearman's correlation coefficient (ρ) for virome targets b), and sex-adjusted quadratic age regression coefficient for virulence factors c). The y-axis represents statistical significance as −log10(q value), with the dotted line indicating the FDR significance threshold ($q < 0.05$). Significant associations are highlighted in green, and labeled targets represent the most significant age-associated features. Directionality of age effects is indicated by the corresponding x-axis orientation. d) Summary of the number of significant age-associated targets identified within each antigen space after FDR correction ($q < 0.05$). Bars indicate the number of significant targets detected in the self-antigen, virome, and virulence factor libraries.

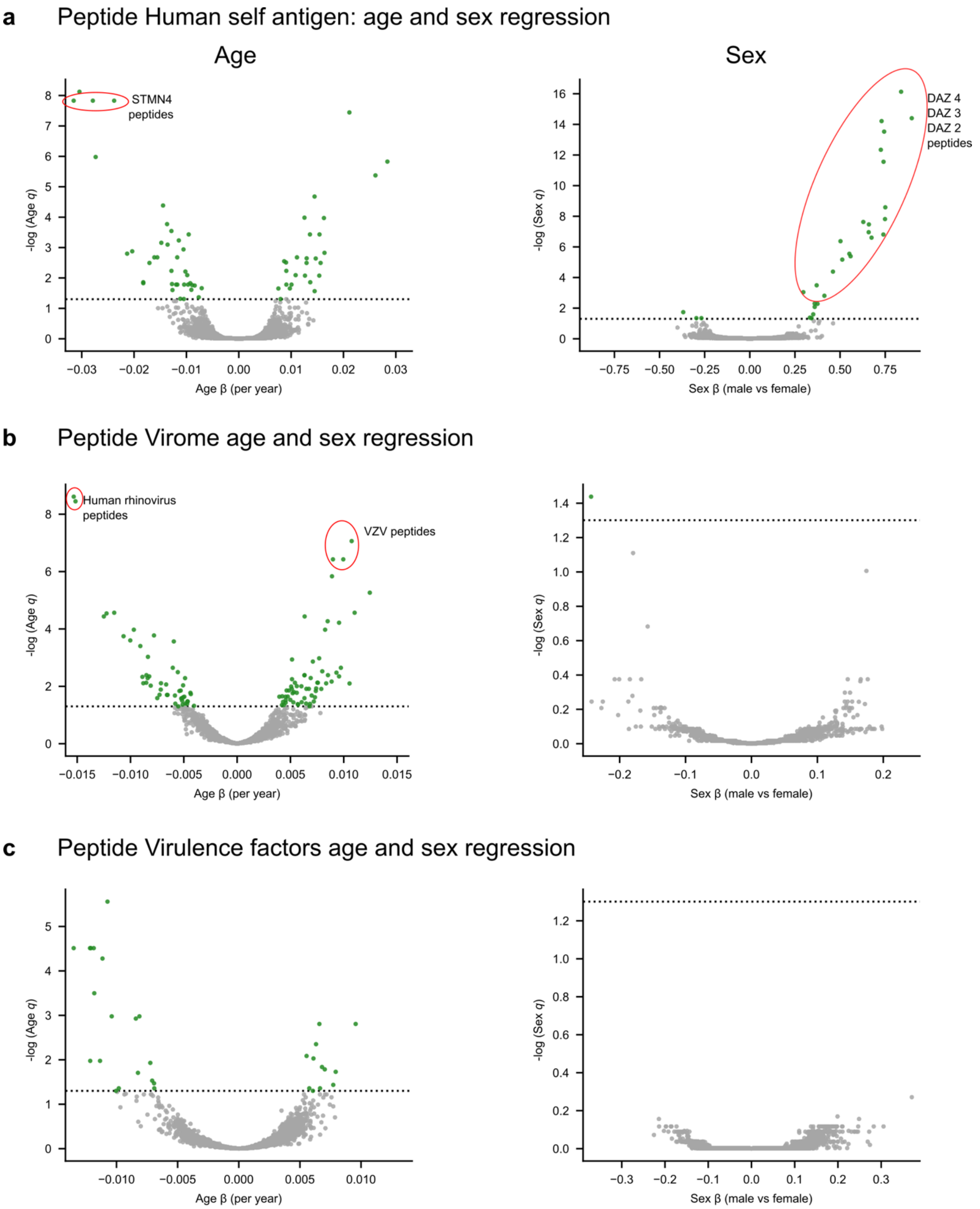


Figure S5. Peptide-level age and sex associations across antigenic libraries.

a) Human self-antigen peptide associations with age and sex. Volcano plots show peptide-level regression coefficients (β) and statistical significance (−log10(q)) for age (left) and sex (male versus female; right). Age models identified significant associations including peptides derived from STMN4 ($q = 1.48 \times 10^{-8}$), CDC73 ($q = 7.49 \times 10^{-9}$), RIN3 ($q = 6.97 \times 10^{-4}$), and QTRT1 ($q = 1.49 \times 10^{-3}$), whereas sex-associated peptides included DAZ2 and DAZ4 (q values as low as $3.95 \times 10^{-15}$ and $4.24 \times 10^{-7}$, respectively). b) Virome peptide associations with age and sex. Peptide-level regression results are shown for viral antigen targets, with age effects quantified by regression coefficients and sex effects by male-versus-female coefficients. Representative age-associated viral targets included peptides from Human rhinovirus 1B ($q = 2.45 \times 10^{-9}$), human rhinovirus 23 ($q = 3.55 \times 10^{-9}$), and varicella-zoster virus ($q = 8.65 \times 10^{-8}$). No broad enrichment of sex-associated virome peptides was observed after FDR correction. c) Virulence factor peptide associations with age and sex. Peptide-level regression analyses across virulence factor antigens identified age-associated targets including peptides from *Staphylococcus haemolyticus* ($q = 3.0 \times 10^{-6}$), *Staphylococcus aureus* ($q = 3.1 \times 10^{-5}$), and *Staphylococcus epidermidis* ($q = 3.1 \times 10^{-5}$). No virulence factor peptides passed the FDR threshold for sex-associated effects.

For all panels, peptide-level associations were evaluated using linear regression models for all peptides positive in at least 5% of samples, with age effects estimated as continuous age coefficients and sex effects estimated as male-versus-female coefficients. P values were adjusted for multiple testing within each antigen library using the Benjamini–Hochberg procedure, and peptides meeting an FDR threshold of $q < 0.05$ were considered significant (green).

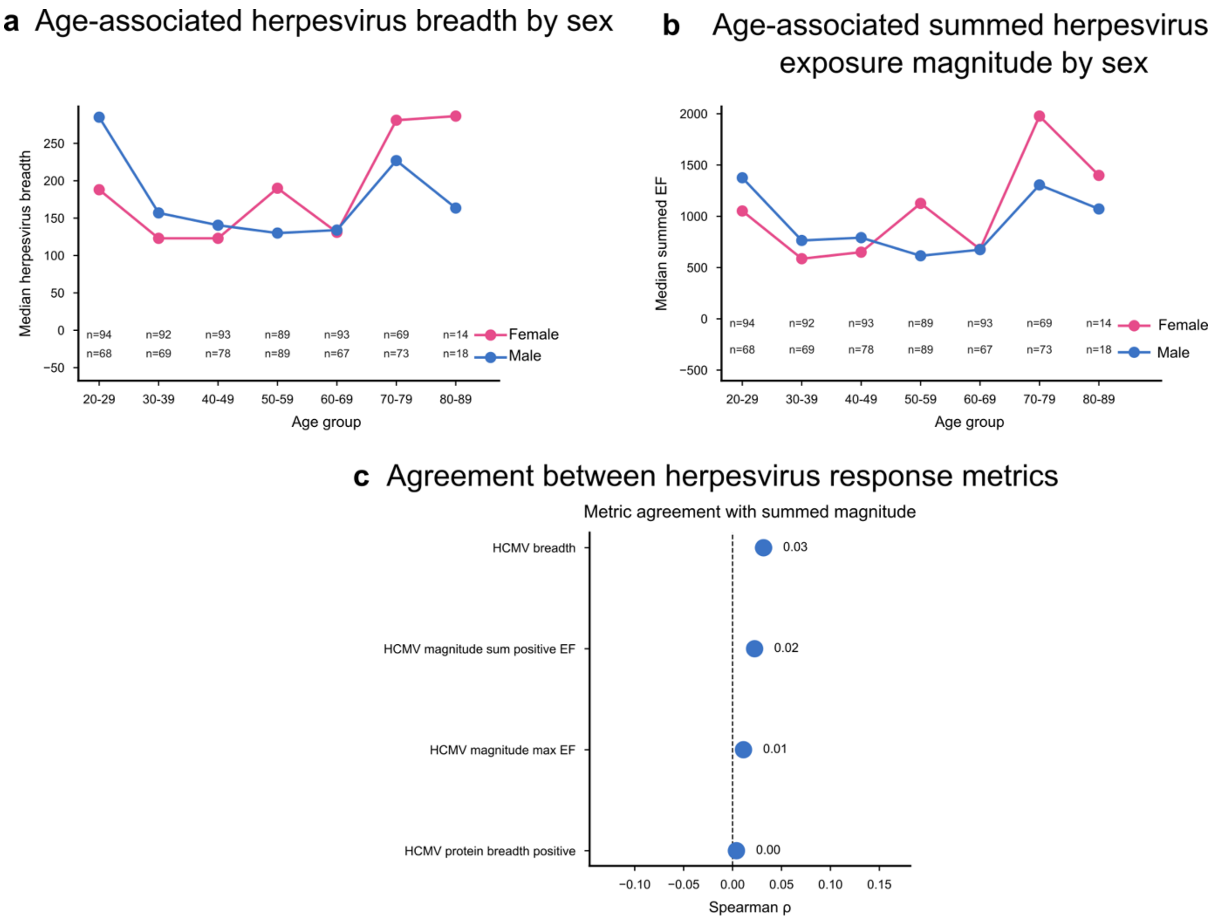


Figure S6. Age- and sex-associated variation in herpesvirus antibody response breadth and magnitude.

a) Age-associated herpesvirus response breadth stratified by sex. Lines show the median number of herpesvirus targets recognized across age groups for females and males; sample sizes are indicated below each age category. b) Age-associated summed herpesvirus exposure magnitude stratified by sex. Lines show the median summed enrichment factor (EF) across herpesvirus responses within each age and sex stratum. c) Agreement between complementary HCMV response metrics. Spearman correlation coefficients between summed response magnitude and alternative response metrics are shown, including HCMV response breadth and magnitude-based measures. Points indicate the correlation strength between metrics, with values labeled.

For all panels, herpesvirus response metrics were summarized at the participant level using PhIP-seq-derived enrichment factors. Age- and sex-stratified comparisons were performed using predefined age bins and sex groups, with sample sizes shown for each comparison. Associations between response metrics were evaluated using Spearman correlation, and effect-size comparisons were assessed using permutation-based statistical testing.

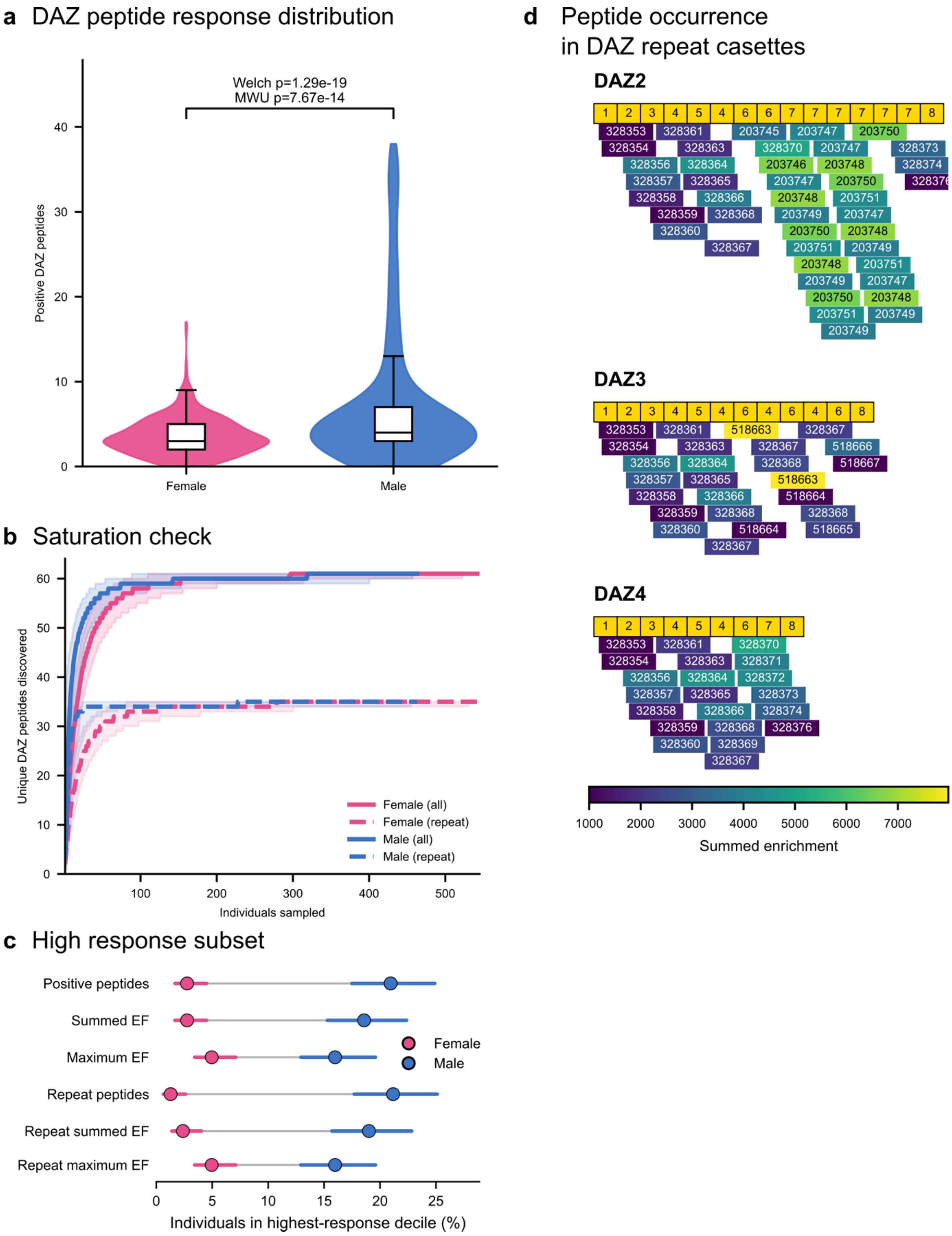


Figure S7. Sex differences and repeat architecture of DAZ peptide responses.

a) Distribution of DAZ peptide response breadth between females and males. Violin plots show the number of DAZ peptides with positive enrichment factor (EF) responses per participant. Males showed significantly higher DAZ peptide breadth than females (Welch's t-test, $p = 1.29 \times 10^{-19}$; Mann–Whitney U test, $p = 7.67 \times 10^{-14}$; Cohen's $d = 0.64$). b) Saturation analysis of DAZ peptide discovery. Curves show the accumulation of unique DAZ peptides identified as increasing numbers of individuals are sampled, stratified by sex and including all DAZ peptides or repeat-associated peptides. Shaded regions indicate 95% confidence intervals. Saturation was assessed to evaluate whether additional sampling increased discovery of unique DAZ peptide responses. c) Sex distribution within the high-response DAZ subset. Cleveland plots show the proportion of females and males represented among individuals in the highest-response decile for multiple DAZ response metrics, including total positive peptides, summed EF, maximum EF, and repeat-associated response metrics. Points represent group proportions with 95% confidence intervals calculated using the Wilson method. d) Occurrence and enrichment of DAZ repeat-rich associated peptides across DAZ repeat cassettes. Peptides are mapped to canonical DAZ repeat regions, with color intensity representing summed enrichment factor across responding individuals. The repeat architecture and peptide localization demonstrate recurrent targeting of specific DAZ repeat-derived regions.

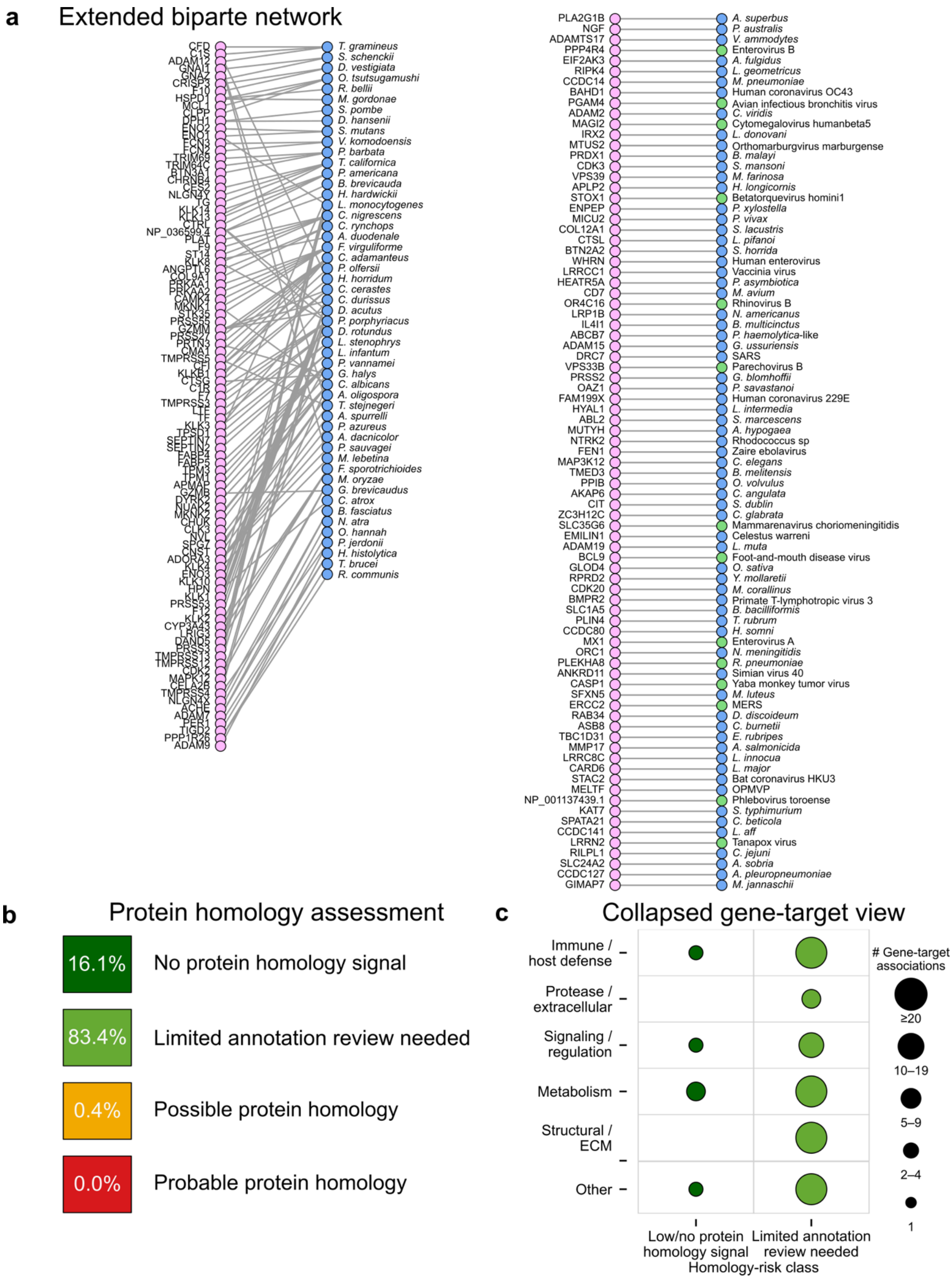


Figure S8. Characterization of cross-library bridge pairs and protein homology assessment.

a) Extended bipartite network of antigen bridge pairs linking human proteins (pink nodes) to microbial, viral, or other non-human targets (blue nodes). Edges represent shared peptide-level associations between antigen targets, identifying bridge pairs with potential sequence or immunological overlap across antigen spaces. Green-highlighted nodes indicate selected targets with reviewed protein homology or annotation evidence. b) Protein homology assessment of bridge-pair targets. Bridge pairs were categorized according to homology confidence based on sequence similarity and annotation review. The majority of pairs showed limited annotation requiring further review (83.4%), whereas smaller fractions showed no detectable protein homology signal (16.1%) or evidence of possible (0.4%) or probable (0.0%) protein homology. c) Collapsed functional overview of bridge-pair gene–target associations. Associations are grouped by broad functional categories and stratified according to protein homology status. Bubble size indicates the number of gene–target associations within each category, highlighting the distribution of bridge-pair annotations across functional classes.

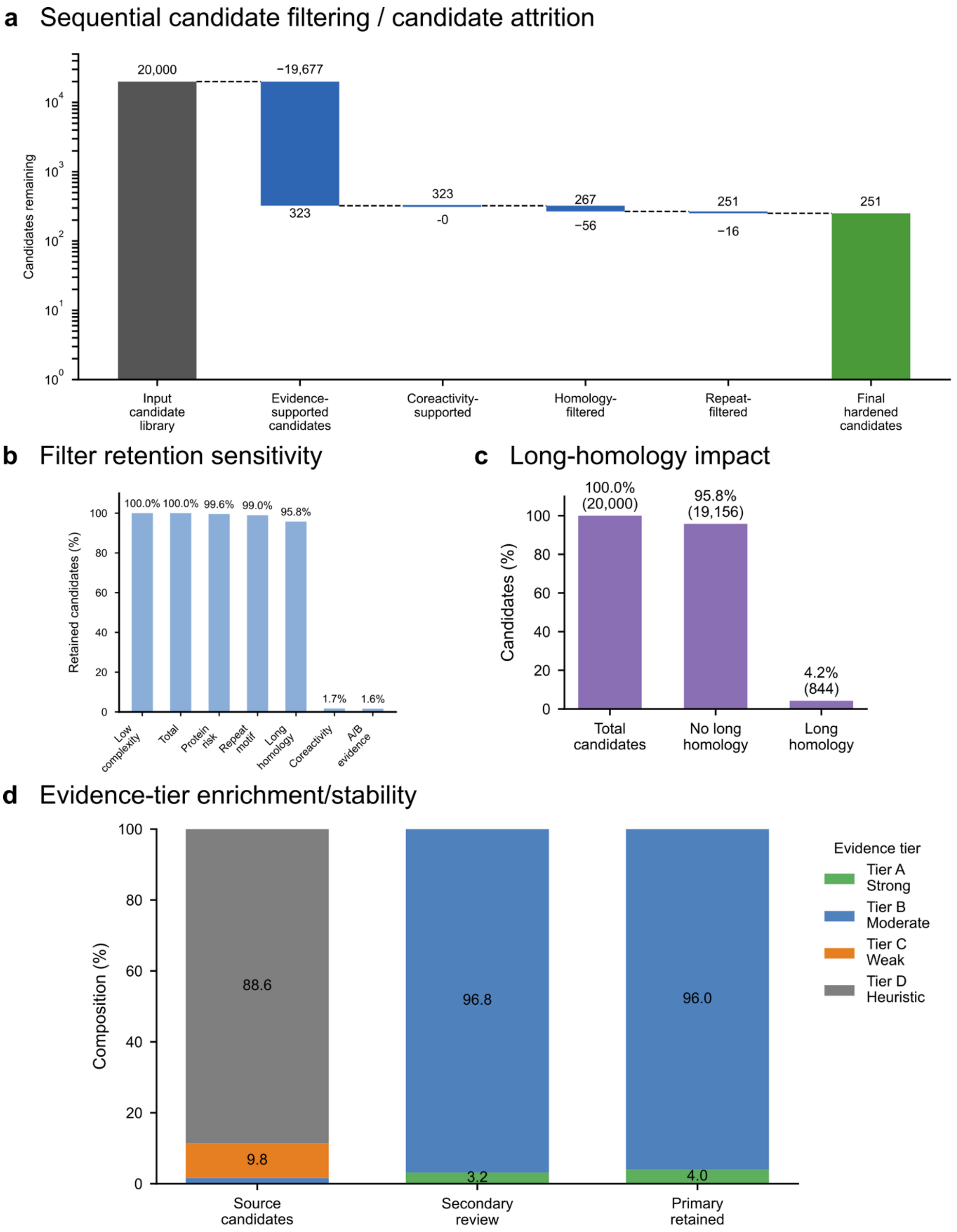


Figure S9. Sequential filtering, sensitivity analysis, and evidence-tier evaluation of prioritized candidates.

a) Sequential candidate filtering workflow showing reduction of the initial candidate library through successive evidence-based filtering steps. Starting from 20,000 input candidates,

candidates were retained after evidence support, co-reactivity assessment, homology filtering, and repeat filtering, resulting in 251 final hardened candidates. b) Sensitivity analysis of candidate retention following sequential filtering criteria. The proportion of candidates retained after applying alternative filtering steps is shown, demonstrating the relative impact of individual filters on candidate prioritization. c) Impact of long-homology filtering on candidate retention. The proportion of candidates affected by removal of long homology matches is shown, with the majority of candidates retained after exclusion of long-homology-associated candidates. d) Evidence-tier enrichment and stability during candidate prioritization. Candidate composition is shown across evidence tiers for the source candidate set, secondary review stage, and final retained candidates. Evidence tiers represent decreasing levels of confidence from Tier A (strong evidence) to Tier D (heuristic evidence), demonstrating enrichment of higher-confidence candidates after review and filtering.

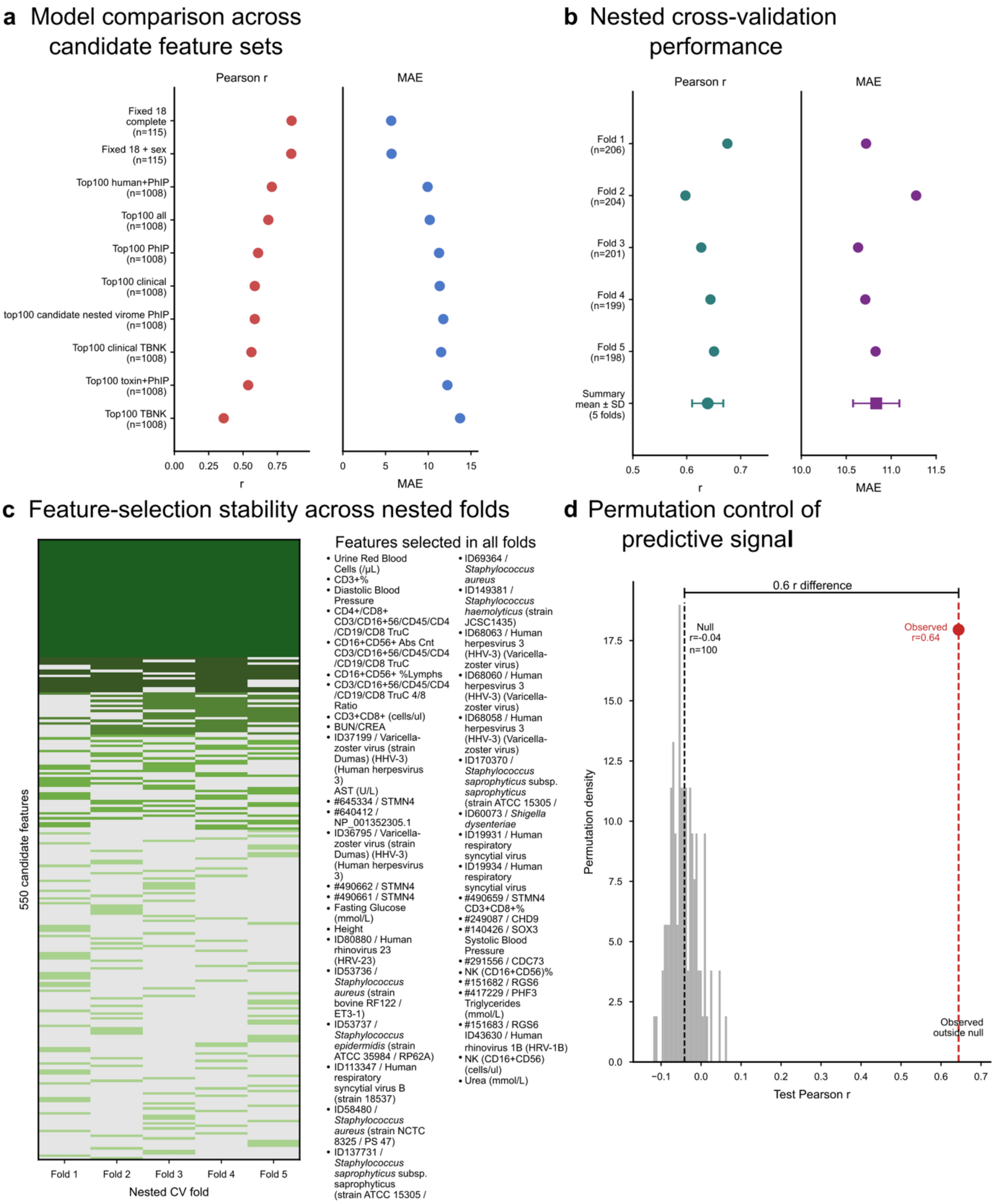


Figure S10. Evaluation of predictive model performance, feature stability, and permutation-based control of predictive signal.

a) Comparison of model performance across candidate feature sets. Predictive performance is shown for alternative feature combinations using Pearson correlation (r) between predicted and observed values and mean absolute error (MAE). Models incorporating broader multimodal feature sets showed improved predictive performance compared with restricted candidate feature sets. b) Nested cross-validation performance assessment. Pearson correlation and MAE are shown across five independent outer folds, with summary values representing the mean ± SD across folds. Consistent performance across folds demonstrates model stability and generalizability. c) Feature-selection stability across nested cross-validation folds. The heatmap shows the selection frequency of candidate features across five validation folds. Features selected in all folds are listed alongside the stability map, highlighting consistently retained predictive features across resampling iterations. d) Permutation-based evaluation of predictive signal. The distribution of Pearson correlations obtained after permutation of outcome labels is compared with the observed model performance. The observed correlation ($r = 0.64$) was substantially greater than the permutation-derived null distribution (mean $r = -0.04$; $n = 100$ permutations), supporting the presence of non-random predictive signal.

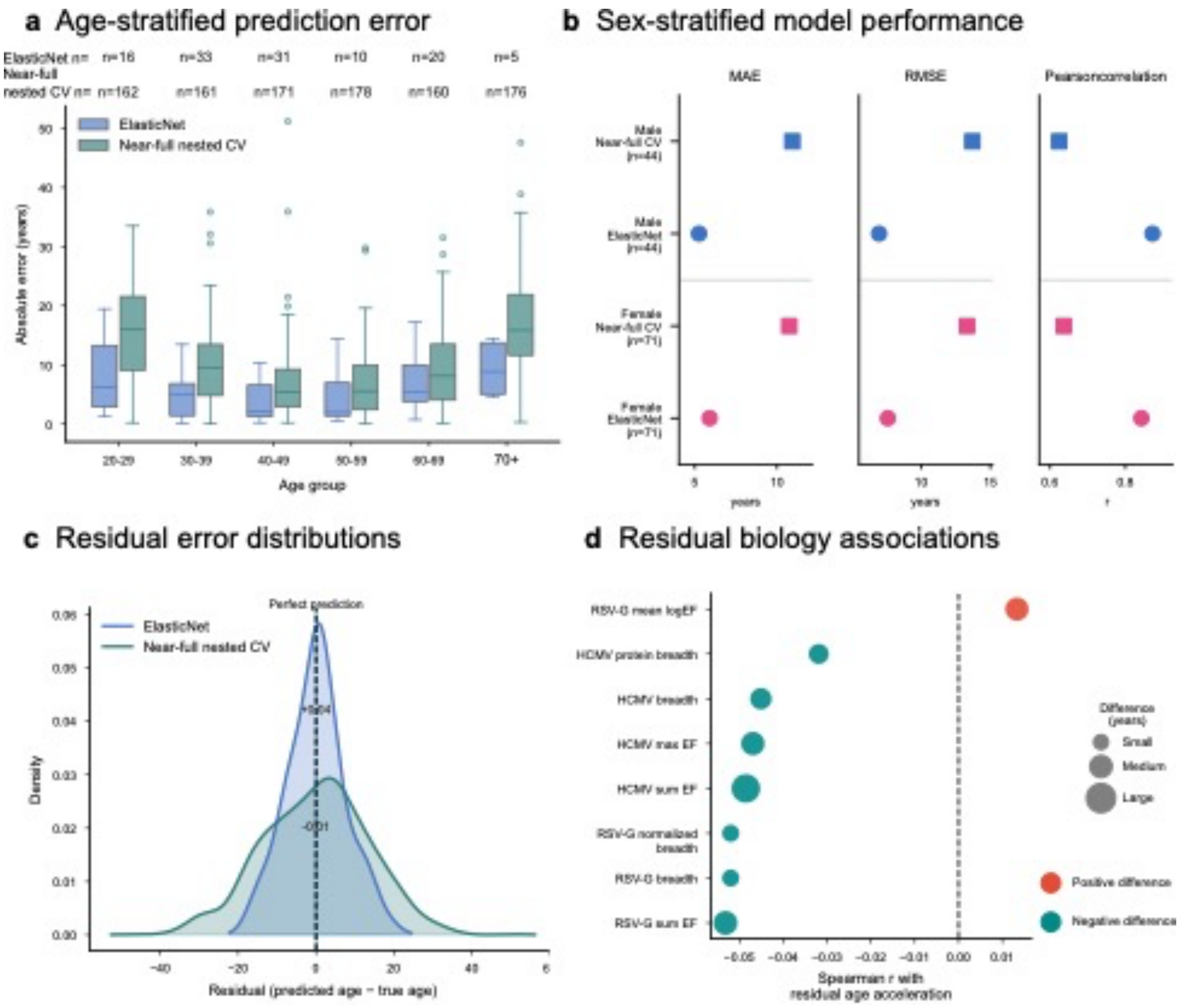


Figure S11. Evaluation of age-prediction performance across demographic strata and residual biological associations.

a) Age-stratified prediction error across age groups. Boxplots show absolute prediction error (years) for ElasticNet and near-full nested cross-validation models across age strata. Sample sizes for each age group are indicated above the distributions. b) Sex-stratified model performance. Model performance metrics are shown separately for females and males using mean absolute error (MAE), root mean squared error (RMSE), and Pearson correlation between predicted and chronological age for ElasticNet and near-full nested cross-validation models. c) Residual error distributions. Density distributions of prediction residuals (predicted age − chronological age) are shown for ElasticNet and near-full nested cross-validation models. The dashed line indicates perfect prediction (residual = 0), with median residual offsets shown for each model. d) Biological associations with age-prediction residuals. Spearman correlations between prediction residuals and selected immune and microbial response features are shown. Point size indicates the magnitude of the associated age difference, and color indicates the direction of association, with positive values representing features associated with overprediction and negative values representing features associated with underprediction.

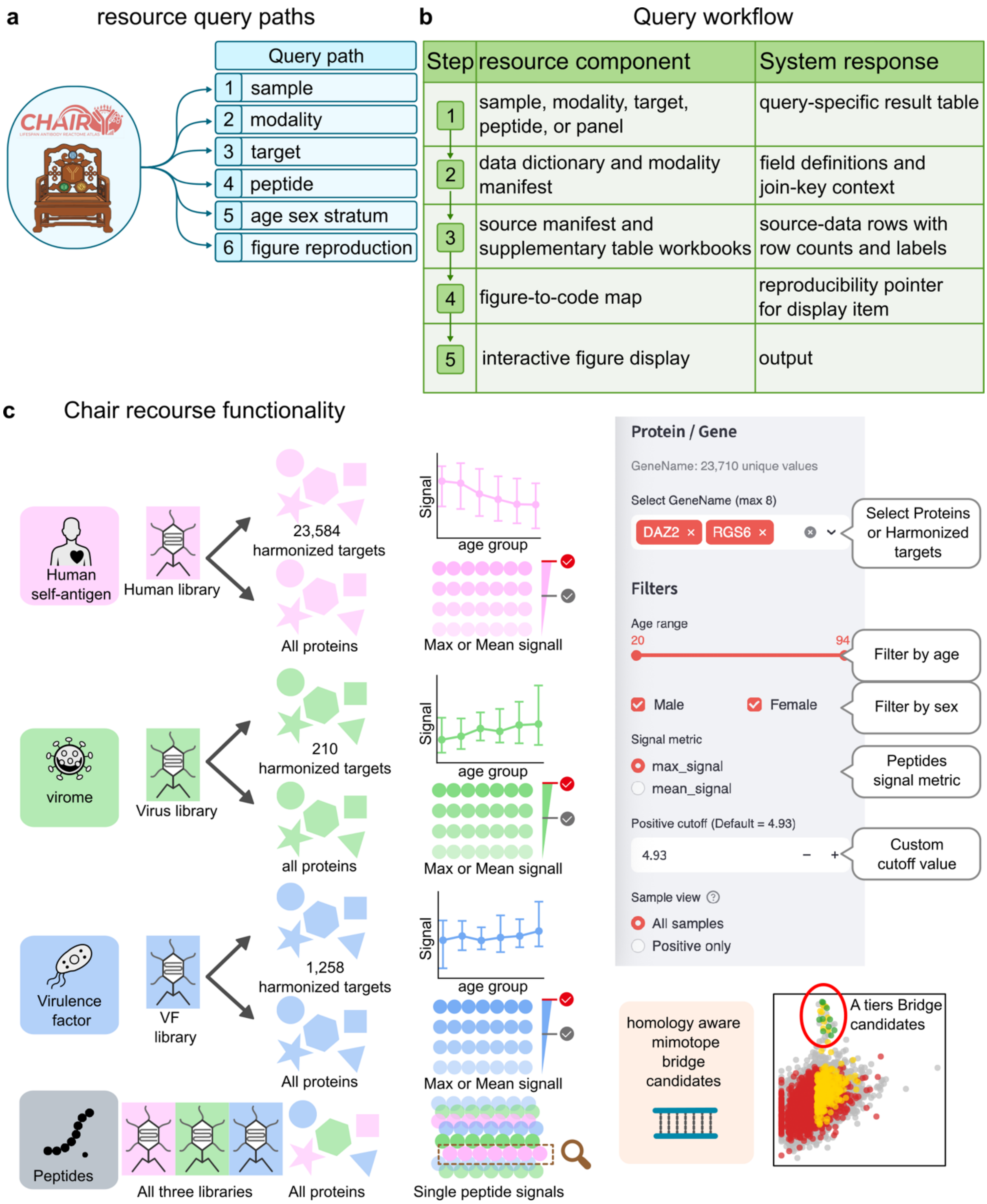


Figure S12. Query architecture and functionality of the CHAIR resource interface.

a) Available resource query paths enabling exploration of CHAIR data by sample, modality, target, peptide, age/sex stratum, and figure reproduction. b) Query workflow illustrating the progression from user-defined queries through resource components, including sample and modality metadata, data dictionaries, source manifests, figure-to-code mappings, and claimboundary fields. Each query step returns context-specific outputs designed to support reproducible data exploration and interpretation. c) CHAIR resource functionality overview. The interface enables interrogation across multiple antigenic libraries, including human self-antigens, viral proteins, and virulence factors, as well as combined library searches. Users can select protein or organism targets, filter samples by age and sex, define peptide signal metrics and thresholds, and visualize peptide-level or protein-level response patterns across the cohort.
